# Elucidating the antiviral mechanism of different MARCH factors

**DOI:** 10.1101/2020.12.22.422953

**Authors:** Supawadee Umthong, Brian Lynch, Uddhav Timilsina, Brandon Waxman, Emily B. Ivey, Spyridon Stavrou

**Author notes:** Address correspondence to Spyridon Stavrou.

## Abstract

The Membrane Associated RING-CH (MARCH) proteins belong to a family of E3 ubiquitin ligases, whose main function is to remove transmembrane proteins from the plasma membrane. Recent work has shown that the human MARCH1, 2 and 8 are antiretroviral factors that target the Human Immunodeficiency virus-1 (HIV-1) envelope glycoproteins by reducing their incorporation in the budding virions. Nevertheless, the dearth of information regarding the antiviral mechanism of this family of proteins necessitates further examination. In this study, using both the human MARCH proteins and their mouse homologues, we provide a comprehensive analysis of the antiretroviral mechanism of this family of proteins. Moreover, we show that human MARCH proteins restrict to varying degrees the envelope glycoproteins of a diverse number of viruses. This report sheds light on the important antiviral function of MARCH proteins and their significance in cell intrinsic immunity.

## Introduction

The viral envelope is critical for viral replication and spread and thus a possible target for host antiviral factors. Such factors include the members of the Membrane Associated RING-CH (MARCH) protein family, a unique family of E3 protein ligases, which target a number of immune receptors found in the plasma membrane (PM)(1). The human and mouse MARCH families consist of 11 members, of which 9 are transmembrane (TM) proteins with at least 2 TM domains (1). MARCH proteins have important regulatory roles in lymphocyte development (2-5) by targeting and removing from the PM a number of cellular membrane proteins including Major Histocompatibility Complex II (MHC II) by MARCH1 and MARCH8 (4, 6-10) and DLG1 by MARCH2 (11). MARCH1 with MARCH8 and MARCH2 with MARCH3 are thought to form two homologous pairs, respectively, due to their structural homology, which suggests that they share similar substrates (1, 12, 13). MARCH1, 2 and 8 are expressed in the spleens, lymph nodes and bone marrow (1, 2, 9, 13), tissues targeted by retroviruses, such as Murine Leukemia virus (MLV) and Human Immunodeficiency virus-1 (HIV-1).

A common feature among all members of the MARCH protein family is the presence of a RING-CH (Really Interesting New Gene-CH) domain, which is a variant RING domain (4) and is critical for the ubiquitination and downregulation of their target proteins (12). Most MARCH proteins contain two or more TM domains that are essential for their localization and for MARCH-mediated target protein recognition (14-17). Additional domains have been identified in MARCH1 and MARCH8 to be important for their function including a <u>d</u>omain in between the <u>R</u>ING-CH domain and the transmembrane domain (DIRT domain), a short C’ terminal sequence (VQNC), which is important for the spatial organization of MARCH1 and MARCH8, and two C’ terminal cytoplasmic tyrosine endocytic motifs (YXXΦ) (15, 18-22).

Immune receptors on the PM targeted by MARCH proteins are degraded either in the lysosome or the proteasome. MHC II, a target of MARCH1 and MARCH8, is degraded in the lysosome (4, 6, 23), while IL-1 receptor accessory protein (IL-1RAcP) is directed by MARCH8 for proteasomal degradation (24). Furthermore, the cytosolic tail of immune receptors (MHC II, CD86) targeted by MARCH proteins is critical for MARCH-mediated degradation (6, 19, 25, 26).

The retroviral Envelope (*env)* gene encodes a surface-exposed glycoprotein that is essential for virus entry into a new cell (27). The extracellular subunit is referred to as the surface (SU) subunit (gp120 for HIV-1 and gp70 for MLV) and is responsible for receptor binding while the TM subunit (gp41 for HIV-1 and p15E for MLV) is important for cell-virus fusion. The TM subunit of MLV gets further cleaved by the viral protease during or shortly after assembly producing the p12E form of TM found in mature virions (28, 29). Recent reports identified human MARCH1, 2 and 8 as HIV-1 restriction factors, which reduce HIV-1 infectivity by blocking the incorporation of gp120 and gp41 in the viral envelope of newly synthesized virions by sequestering them intracellularly (30-32). Nevertheless, little is known about their mechanism of restriction, how much conserved or how broad is their antiviral function.

In this report, using both HIV-1 and MLV as well as human and mouse MARCH proteins, we found that unlike previous reports, HIV-1 and MLV envelope glycoproteins are not sequestered intracellularly but are degraded in the lysosome. In addition, we identified the domains of mouse MARCH1 and 8 that are important for restriction and for physically binding to the retroviral envelope glycoproteins. Furthermore, using a variety of viral envelope glycoproteins, we demonstrated that human MARCH proteins have broad antiviral functions and target for degradation a variety of viral glycoproteins including those of Lymphocytic Choriomeningitis Virus (LCMV), Lassa Virus (LASV) and Severe Acute Respiratory Syndrome Coronavirus 2 (SARS-CoV-2).

## Results

### Transcriptional regulation of mouse MARCH proteins

Human MARCH1 and MARCH2, but not human MARCH8, are type I interferon (IFN) inducible genes (32). Thus, we examined the effect of IFN-β on the murine homologues of MARCH1, 2 and 8 in different primary and stable cell lines. We treated bone marrow derived dendritic cells (BMDCs), bone marrow derived macrophages (BMDMs), EL4 (murine T lymphocyte cell line), NIH3T3 (murine fibroblast cell line) and MutuDC1940 (immortalized mouse dendritic cell (DC) cell line (33)) with IFN-β (500u/ml). We collected RNA at different time points and performed RT-PCR to determine mouse MARCH1, 2, 3 and 8 expression levels. We found that mouse MARCH1 was expressed only in MutuDC1940, BMDCs and BMDMs while mouse MARCH2 and MARCH8 were expressed in all cell lines tested (Fig. 1A to C and S1A and B). Mouse MARCH3 was expressed only in stable cell lines (Fig. 1C, S1A to C) and not in primary cells (Fig. 1A and B). Interestingly, only mouse MARCH1 was IFN-β inducible in all cell lines expressing it (BMDMs, BMDCs, and MutuDC1940) (Fig. 1A to C). Mouse MARCH3, 8 and 2, unlike human MARCH2, were not IFN-β inducible (Fig. 1A to C and S1A and B). By treating MutuDC1940 with varying amounts of IFN-β, we found that IFN-β upregulated mouse MARCH1 mRNA levels in a dose-dependent manner (Fig. 1D). To determine the effect of MLV infection on the expression levels of mouse MARCH1, 2, 3 and 8, we infected MutuDC1940, EL4 and NIH 3T3 with MLV (5 MOI) and at different time points collected RNA and performed RT-PCR to determine their expression levels. We found that MLV infection had no effect on mouse MARCH1, 2, 3 and 8 expression in all cell lines tested (Fig. 1E, F and S1C). Thus, we concluded that MLV infection has no effect on mouse MARCH1, 2, 3 and 8 transcript levels and only mouse MARCH1 is an IFN stimulated gene.

**Fig 1.**
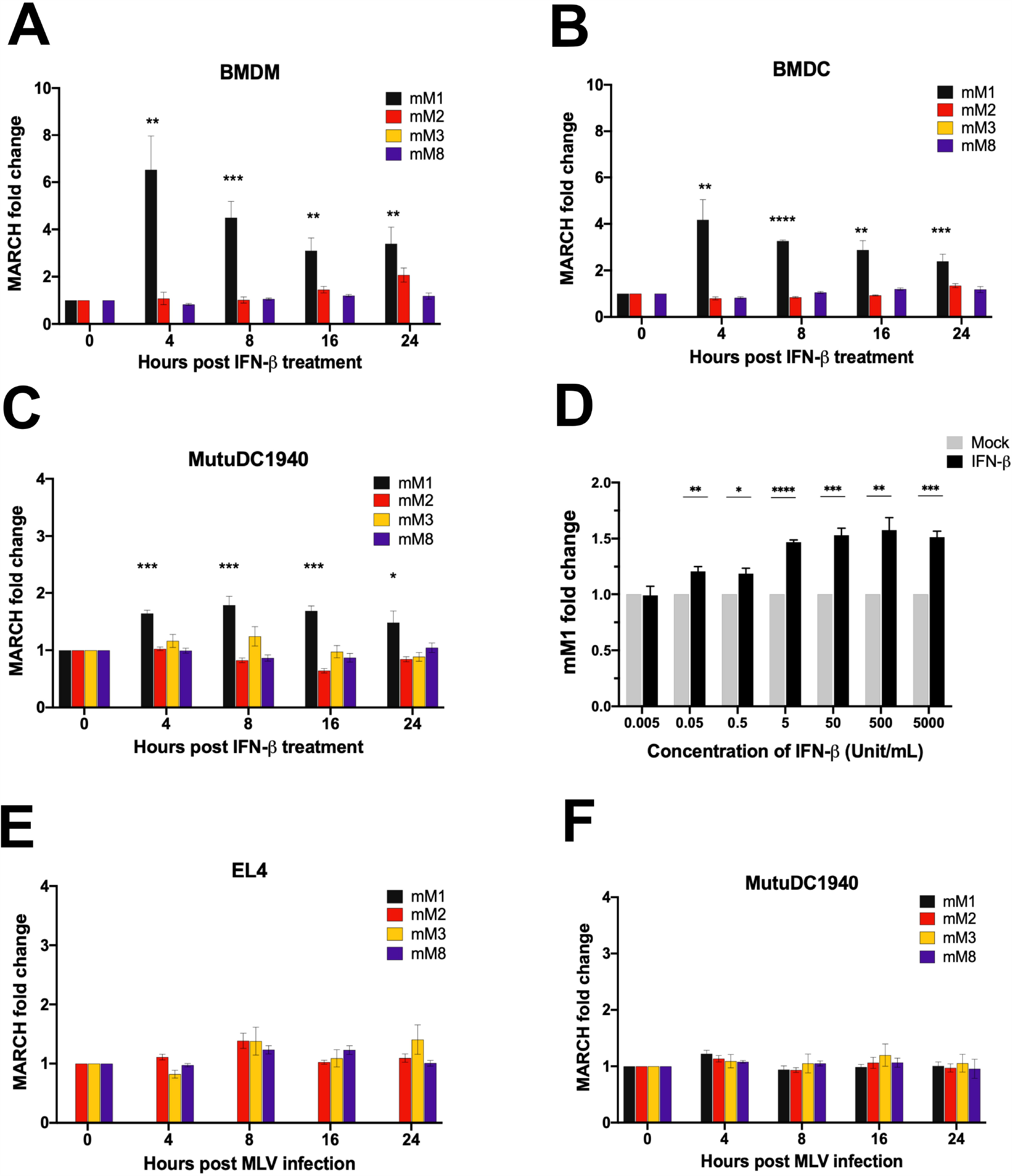
Expression levels of mMARCH1, 2, 3, and 8 in the presence of IFN-β and MLV infection. Fold expression changes of mouse MARCH1, 2, 3 and 8 (mM1, 2, 3 and 8) relative to untreated cells and normalized to GAPDH in (A) bone marrow derived macrophages (BMDMs), (B) bone marrow derived dendritic cells (BMDCs) and (C) MutuDC1940 cells at 0, 4, 8, 16 and 24 hours post treatment with 500 Units/mL of mouse IFN-β. Purity of BMDMs and BMDCs was determined by flow cytometry staining with anti-CD11b and anti-CD11c respectively as shown in Fig. S1D and S1E in the supplementary material. (D) Fold expression changes of mM1 relative to untreated cells and normalized to GAPDH after treatment of MutuDC1940 cells with different concentrations of IFN-β (0.005 to 5000 Units/mL) for 4 hours. (E-F) Fold expression changes of mM1, 2, 3, and 8 relative to uninfected cells and normalized to GAPDH in (E) mouse T lymphocyte cell line EL-4 and (F) MutuDC1940 cells at 4, 8, 16 and 24 hours post infection with MLV (5 MOI). Data shown represent averages of results from n=3 independent experiments. All results are presented as means ± standard error of the mean (SEM). Statistical analysis performed using unpaired t-test. * = p ≤ 0.05, ** = p ≤ 0.01, *** = p ≤ 0.001, **** = p ≤ 0.001.

### Mouse MARCH1 and 8 do not sequester the MLV and MMTV envelope glycoproteins intracellularly but target them for degradation

To determine the effect mouse MARCH genes have on retrovirus envelope glycoproteins, we co-transfected 293T cells with an MLV molecular clone and with either an empty vector (E.V.) or mouse MARCH1, 2, 3 or 8. Cells and the media were harvested 48 hours post-transfection followed by western blots to determine the levels of the MLV envelope glycoproteins (gp70/SU and p15E/p12E/TM). In the cell fractions, we found that the protein levels of both gp70 and p15E were reduced in the presence of mouse MARCH1 and mouse MARCH8, while mouse MARCH2 and 3 had no effect (Fig. 2A). Furthermore, virions produced in the presence of mouse MARCH1 or 8 had significantly lower levels of gp70 and p12E than virions produced in cells transfected with either E.V., mouse MARCH2 or 3 (Fig. 2A). By using varying levels of mouse MARCH1 and mouse MARCH8, we found that mouse MARCH1 and 8 degraded gp70 and p15E in a dose dependent manner (Fig. 2B). Interestingly, when probing for p15E in the cell fractions, we observed a band (shown with an asterisk) migrating slightly faster than p15E when using higher amounts of either mouse MARCH1 or mouse MARCH8, which probably reflects a degradation intermediate of p15E (Fig. 2B). To determine if the decreased incorporation of gp70 and p15E in the nascent virions affected virus infectivity, we produced MLV-luciferase reporter viruses in the presence of either mouse MARCH1, 2, 3, 8 or E.V. NIH3T3 cells were infected with equal amounts of MLV CA and luminescence was measured 48 hours post-infection. We found that virions produced in the presence of mouse MARCH1 or 8 were significantly less infectious compared to those produced in the presence of mouse MARCH2 or 3 (Fig. 2C). We also investigated the effect of mouse MARCH1, 2, 3 and 8 on the viral envelope glycoproteins of another murine retrovirus, mouse mammary tumor virus (MMTV). We co-transfected 293T cells with a plasmid encoding an infectious genetically engineered MMTV hybrid provirus (HP) (34) and either mouse MARCH1, 2, 3, 8 or E.V. Cell and media fractions were harvested and western blots were performed to determine the levels of the MMTV envelope glycoproteins (gp52/SU and gp36/TM). Interestingly, while mouse MARCH1 and mouse MARCH8 abrogated the cellular levels of gp36, they had no effect on the gp52 cellular levels of MMTV (Fig. 2D). Furthermore, mouse MARCH2 lead to reduced levels of only gp36 (Fig. 2D). When examining the virus fraction, we found that virus levels were abrogated in the cells transfected with mouse MARCH1 and mouse MARCH8, while mouse MARCH2 decreased virus production levels (Fig. 2D). This is different from what we observed with MLV, where in the presence of mouse MARCH1 and 8 we detected virions either lacking or with very low levels of incorporated viral envelope glycoproteins (Fig. 2A), similar to what was seen with HIV-1 (30-32). It is possible that the differences in the effect of mouse MARCH1 and 8 on the virus fractions between MLV and MMTV (bald virus vs. no virus) may be attributed to differences in the mechanisms and sites of assembly and release between these two murine retroviruses (35, 36).

**Fig 2.**
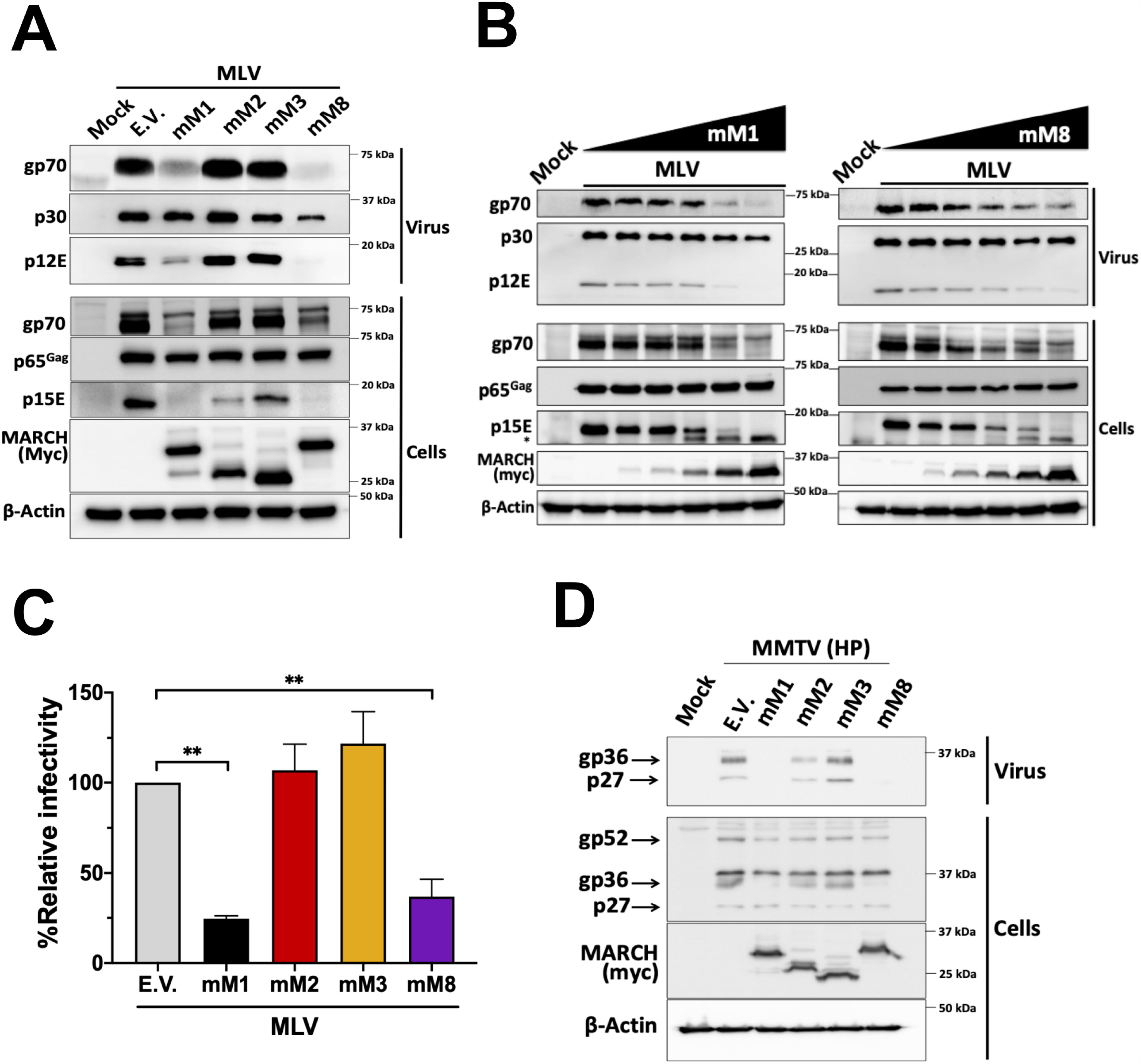
Mouse MARCH1 and mouse MARCH8 block retrovirus infection by targeting the retroviral envelope glycoproteins. (A) Mouse MARCH1 (mM1) and mM8 block the incorporation of the MLV envelope glycoproteins (p15E and gp70) in nascent virions by targeting them for degradation. (B) mM1 and mM8 target the MLV envelope glycoproteins for degradation in a dose dependent manner. For both (A) and (B), 293T cells were co-transfected with an MLV molecular clone and the indicated mouse MARCH constructs. At 48 hours post transfection, cells and released virus in the culture media were harvested and the indicated proteins were analyzed by immunoblotting using anti-MLV p30 (detects both p30 and p65^Gag^), anti-MLV gp70, anti-MLV p15E/p12E, anti-myc (for detection of mM1, 2, 3 and 8) and anti-β-actin antibodies. (C) Virus produced in the presence of mM1 and mM8 has decreased infectivity. NIH3T3 cells were infected with equal amounts of 293T-derived MLV-luciferase reporter virus produced in the presence of mM1, 2, 3, 8 or empty vector (E.V.). Cells were harvested 24 hours post infection and luciferase levels were measured. The percentage (%) of relative infectivity was determined with respect to virus produced in the presence of E.V. All results are presented as means ± SD. Statistical significance was determined by one-way ANOVA. **, p<.01 (D) mM1 and mM8 target MMTV gp36 for degradation. 293T cells were co-transfected with an infectious genetically engineered MMTV hybrid provirus (HP) and either mM1, 2, 3 and 8 or E.V. At 48 hours post transfection, cells and released virus in the culture media were harvested and the indicated proteins were analyzed by immunoblotting using anti-MMTV, anti-myc (for detection of mM1, 2, 3 and 8) and anti-β-actin antibodies. Results are shown for n=3 independent experiments. Representative immunoblotting results are shown in panels A, B and D.

### Human and mouse MARCH proteins differ in their abilities to degrade retroviral envelope glycoproteins

In contrast to previous studies, our aforementioned data show that the cellular levels of the MLV envelope glycoproteins (gp70 and p15E) are reduced in the presence of mouse MARCH1 and 8, suggesting that the viral envelope glycoproteins are degraded and not sequestered intracellularly. We hypothesized that the reason for this discrepancy is due to the difference in the species of origin for the MARCH proteins we used in our system (mouse) vs. in previous reports (human). To investigate this, we co-transfected 293T cells with an MLV molecular clone and with either the human or mouse MARCH1, 2, 8 or an E.V. We observed that both human and mouse MARCH1 and 8 abrogated the cellular levels of gp70 and p15E (Fig. 3A). Interestingly, human MARCH2, unlike mouse MARCH2, significantly reduced the cellular gp70 and p15E levels (Fig 3A). We thus concluded that mouse and human MARCH1 and 8 as well as human MARCH2 degrade and do not sequester intracellularly the MLV envelope glycoproteins.

**Fig 3.**
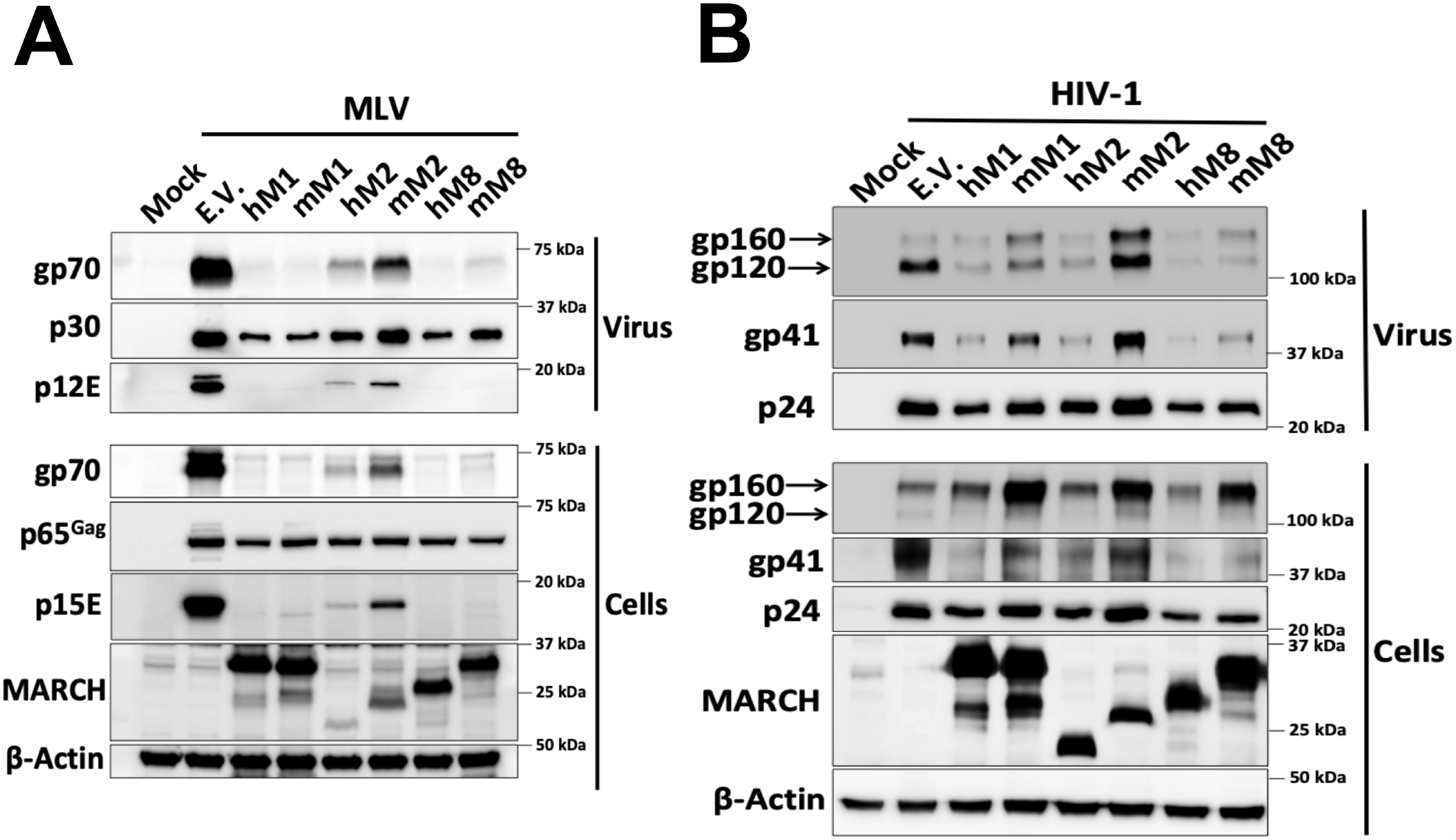
Mouse MARCH and human MARCH proteins differ in their ability to target retroviral envelope glycoproteins. (A) MLV envelope glycoproteins are targeted for degradation by mouse MARCH (mM1) and mM8 and human MARCH (hM1), hM2 and hM8. (B) HIV-1 envelope glycoproteins are targeted for degradation by mM 8 and hM1, 2 and 8. For both (A) and (B), 293T cells were co-transfected with (A) MLV or (B) HIV-1 infectious clone and h/mM1, 2, 8. At 48 hours post transfection, cells and released virus in the culture media were harvested and the indicated proteins were analyzed by immunoblotting using anti-MLV p30 (detects both p30 and p65^Gag^), anti-MLV gp70, anti-MLV p15E/p12E, anti-HIV-1 envelope (detects both gp120 and gp160), anti-gp41, anti-p24, anti-myc (for detection of mM1, 2 and 8 and hM1, 2 and 8) and anti-β-actin antibodies. Results are shown for n=3 independent experiments. Representative immunoblotting results are shown for A and B.

We then speculated that HIV-1 envelope glycoprotein intracellular sequestration by the human MARCH proteins might be virus-specific. Therefore, we examined the effect of human and mouse MARCH1, 2 and 8 on HIV-1. 293T cells were co-transfected with NL4-3 (an HIV-1 lab strain) and either human or mouse MARCH1, 2 and 8 or an E.V. We found that all human MARCH proteins (MARCH1, 2 and 8) decreased the cellular levels of both HIV-1 envelope glycoproteins, gp120 and gp41 (Fig. 3B). On the other hand, only mouse MARCH8 markedly decreased the gp120 and gp41 cellular levels (Fig. 3B). Mouse MARCH2, similar to what we observed with MLV, had no effect (Fig. 3B). Mouse MARCH1, whereas it potently restricted the MLV envelope glycoproteins, had a modest effect on the HIV-1 envelope glycoproteins (Fig. 3B). In summary, our data show that while all human MARCH proteins tested are potent restrictors of retroviral envelope glycoproteins, of the mouse MARCH homologues only mouse MARCH8 was able to potently restrict both MLV and HIV-1 envelope glycoproteins. Finally, in contrast to previous reports, we found that MARCH proteins, both the human and mouse homologues, cause degradation and not intracellular sequestration of the retroviral envelope glycoproteins.

### Mapping the regions of mouse MARCH1 and mouse MARCH8 responsible for their antiviral function

Mouse MARCH1 and 8, due to their homology, have sequence and structural similarities (1, 4, 9). Both mouse and human MARCH1 and 8 have a variant RING domain (RING-CH) that is located in the N’ terminal cytoplasmic tail (Fig. 4A and B) and is essential for their function (1, 30-32), a DIRT domain (18), a VQNC domain (15, 19, 20, 22), two tyrosine (Y) motifs (YXXΦ) present in the C’ terminal cytoplasmic tail (15, 21) and two TM domains (14) (Fig. 4A and B). To determine the importance of these domains on retrovirus restriction, we generated a series of mouse MARCH1 and MARCH8 constructs with mutations or deletions in the various domains. We first verified that the mutant mouse MARCH1 and MARCH8 constructs continued to localize in cellular membranes. In the case of mouse MARCH1, we transfected 293T cells with the wild type and mutant MARCH1 constructs and then stained the cell surface of the transfected cells with an anti-MARCH1 antibody followed by flow cytometry and found that all mouse MARCH1 constructs localize on the plasma membrane (Fig. S2A). Unfortunately, there was no suitable mouse MARCH8 antibody available for flow cytometry. Thus, we transfected 293T cells and isolated membrane bound proteins followed by western blots probing for mouse MARCH8 and for GAPDH to verify the purity of our membrane fractions. We found all mutant MARCH8 constructs localized to cellular membranes similar to wild type MARCH8 (Fig. S2B and C). Subsequently, we co-transfected 293T cells with an MLV molecular clone and the various mouse MARCH1 and mouse MARCH8 mutant constructs followed by western blot to detect any changes in the gp70 and p15E protein levels. We found the RING-CH domain to be essential for MARCH1 and 8-mediated retrovirus restriction (Fig. 4C). Furthermore, deletion of the mouse MARCH8 DIRT domain (ΔDIRT) rescued gp70 and p15E from MARCH8-mediated degradation (Fig. 4D, left panel); however, deletion of the mouse MARCH1 DIRT domain (ΔDIRT) had only a partial effect (Fig. 4D, right panel). Mutations in the VQNC motif (mouse MARCH1/VQNC^mut^ and MARCH8/VQNC^mut^) or the two YXXΦ endocytic motifs had no effect on MARCH-mediated degradation of gp70 and p15E (Fig. 4E and S3). To examine the role of the TM domains, we independently swapped the TM domains of mouse MARCH1 with those of mouse MARCH3, as mouse MARCH3 has no antiretroviral function. We found that only the second TM domain was important for its antiretroviral activity (Fig. 4F, left panel). We initially exchanged the MARCH8 TM domains with those of mouse MARCH3, but it affected protein expression (data not shown). We thus swapped the TM domains of mouse MARCH8 with those of mouse MARCH4, as mouse MARCH4 has no antiretroviral effect (Fig. S4A). We observed that similar to mouse MARCH1, swapping the second TM domain of mouse MARCH8 with that of mouse MARCH4 rendered mouse MARCH8 unable to restrict gp70 and p15E (Fig. 4F right panel). Our findings show that only the RING-CH, DIRT and the second TM domain of mouse MARCH1 and MARCH8 are critical for retroviral envelope glycoprotein restriction.

**Fig 4.**
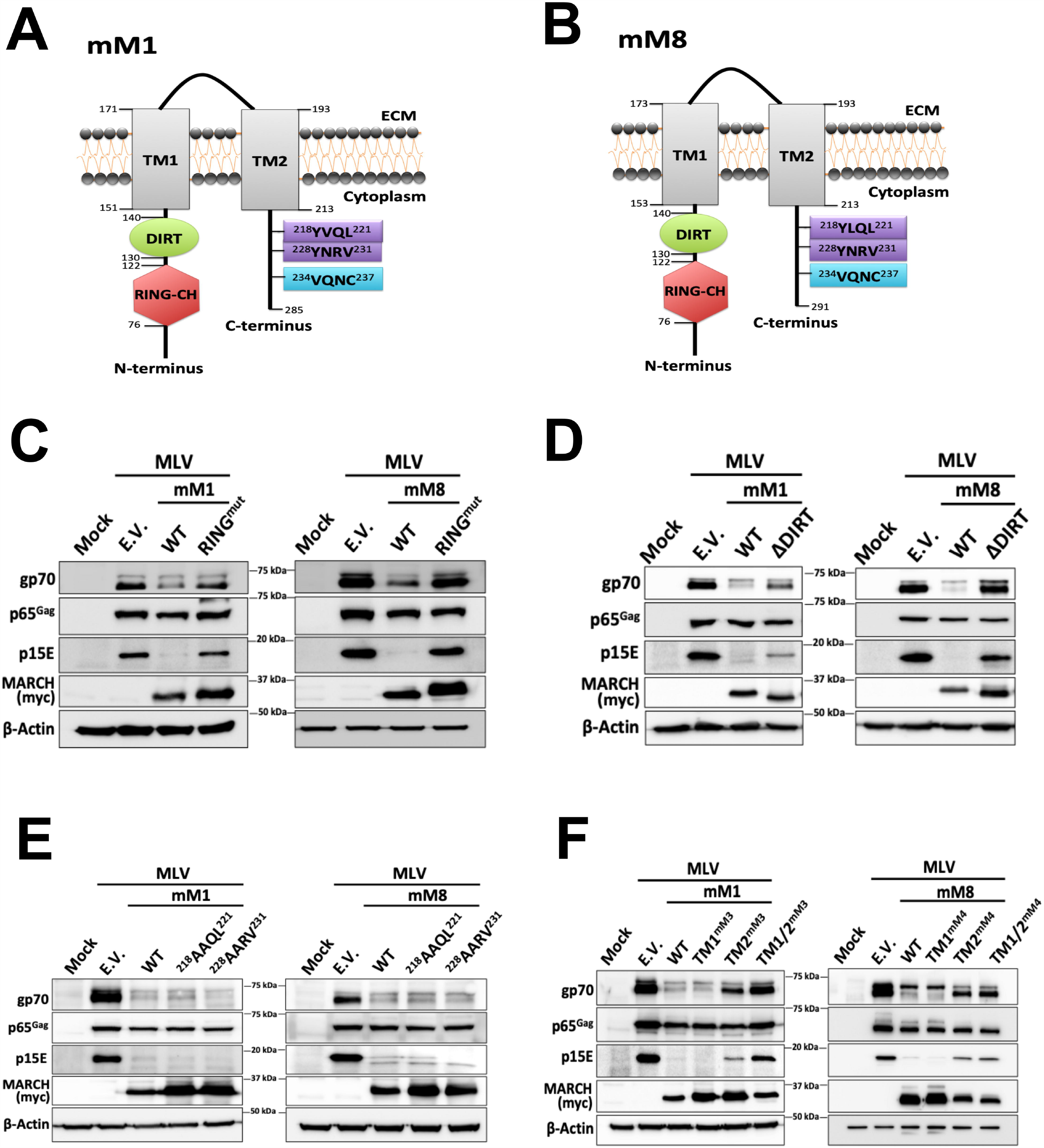
The role of different mouse MARCH1 and mouse MARCH8 domains on retroviral envelope glycoprotein restriction. (A) Schematic diagram of mouse March (mM1) and (B) mM8 and their respective domains: Really Interesting New Gene-CH (RING-CH) domain, the <u>d</u>omain In between <u>R</u>ING-CH domain and <u>T</u>ransmembrane domain (DIRT), N’ terminal and C’ terminal TM domains (TM1 and TM2 respectively), two tyrosine endocytic motifs and a VQNC motif. (C-F) Only the RING-CH, DIRT and TM2 domains of mM1 and 8 are important for the degradation of MLV envelope glycoprotein. 293T cells were co-transfected with either (C) RING-CH mutant (RING^mut^) mM1 and mM8, (D) mM1 and mM8 with the DIRT domain deleted (ΔDIRT), (E) mM1 or mM8 with mutations in the tyrosine endocytic motifs and an MLV infectious clone or (F) mM1 or mM8 with either the N’ terminal (TM1), C’ terminal (TM2) or both TM domains (TM1/2) swapped either with those of mM3 (in the case of mM1) and mM4 (in the case of mM8). Cells were harvested 48 hours post transfection and lysates were analyzed by immunoblotting using anti-MLV p30 (detects p65^Gag^), anti-MLV gp70, anti-MLV p15E, anti-myc (for detection of the mM1 and 8 proteins) and anti-β-actin antibodies. Results are shown for n=3 independent experiments. Representative immunoblotting results are shown for C through F.

### The cytosolic tail of the MLV envelope glycoprotein is crucial for MARCH-mediated restriction

Previous studies have shown that the cytoplasmic tails of the immune receptors targeted by MARCH proteins (e.g. MHC II) are critical for their removal from the cell surface (6, 15, 19, 26, 37). The MLV p15E contains a C’ terminal cytosolic tail that is 35 amino acids long (38). To determine if the cytoplasmic tail of p15E is critical for its MARCH-mediated removal from the PM, we created an MLV molecular clone that did not express the cytoplasmic tail of p15E (pMLVΔCT) by introducing a stop codon at the N’ terminus of the p15E cytoplasmic tail region. We co-transfected cells with an MLV infectious clone expressing either an intact p15E cytoplasmic tail (pMLV) or pMLVΔCT in the presence of either mouse MARCH1, mouse MARCH8 or E.V. We found that p15E with an intact cytoplasmic tail was degraded in the presence of either mouse MARCH1 or 8, while p15E lacking the cytoplasmic tail was resistant to the deleterious effect of mouse MARCH1 and 8 (Fig. 5A). Paradoxically, we repeatedly noticed that p15E lacking the cytoplasmic tail was detected in western blots at higher levels in the presence of either MARCH1 or MARCH8 when compared to E.V. (Fig. 5A), the reason for which is unclear.

**Fig 5.**
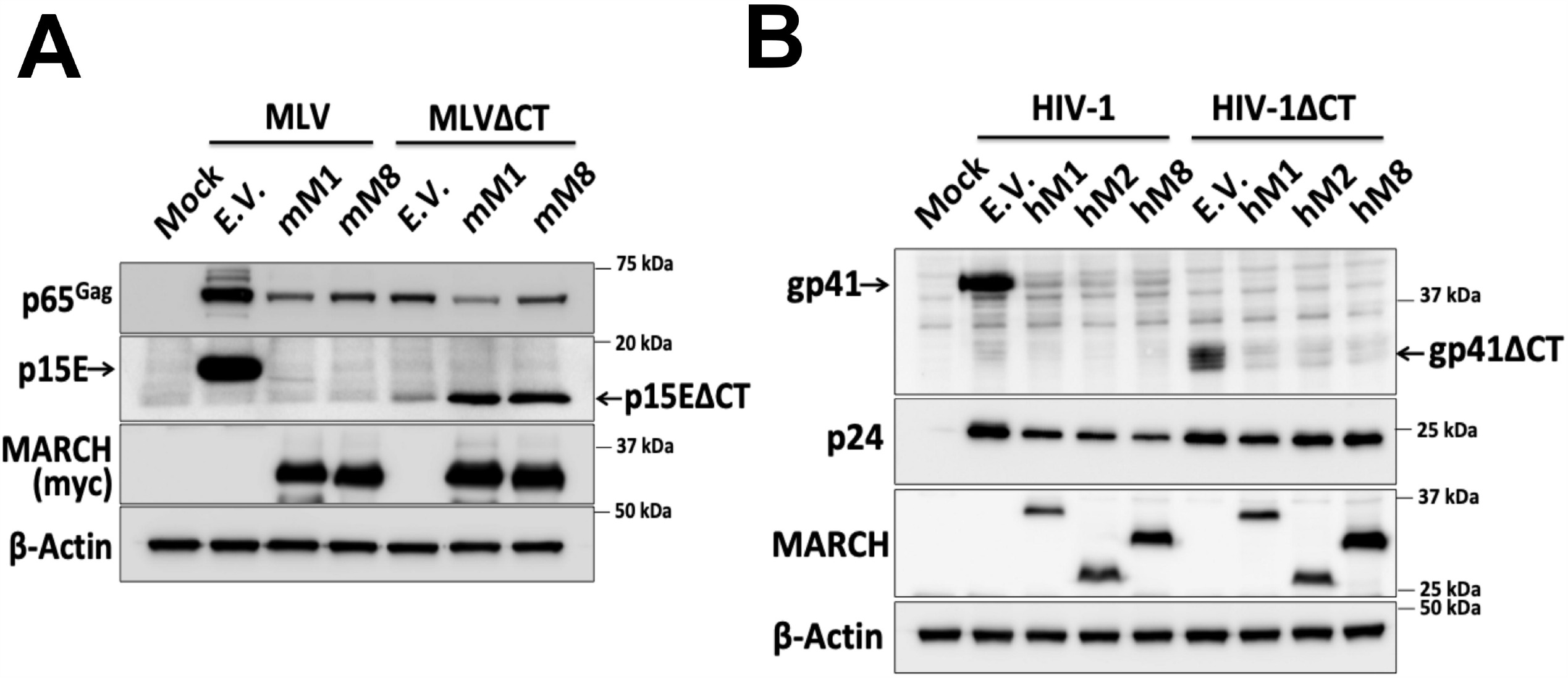
The MLV envelope glycoprotein cytoplasmic tail is critical for MARCH-mediated restriction, but the HIV envelope cytoplasmic tail is dispensable. 293T cells were co-transfected in (A) with wild type MLV or MLV with the cytosolic tail deleted (MLVΔCT) along with either mouse MARCH1 (mM1), mM2, or mM8 and in (B) with wild type HIV or HIV with the cytosolic tail deleted (HIVΔCT) along with either human MARCH (hM1), (hM2) or (hM8). For (A) and (B), 48 hour post transfection cells were harvested and lysates were analyzed by immunoblotting using anti-MLV p30 (detects p65^Gag^), anti-MLV p15E, anti-myc (for detection of human and mouse MARCH proteins), anti-gp41, anti-p24 and anti-β-actin antibodies. Results are shown for n=3 independent experiments. Representative immunoblotting results are shown for A and B.

The retroviral envelope glycoprotein cytoplasmic tail ranges in size, with the cytoplasmic tail of lentiviruses being the longest (27). Due to the differences in size and composition between the MLV p15E and HIV-1 gp41 cytoplasmic tails, it is possible that unlike the MLV p15E cytoplasmic tail, the HIV-1 gp41 cytoplasmic tail is dispensable for MARCH-mediated restriction. Therefore, we generated an HIV-1 molecular clone that does not express the cytoplasmic tail of gp41 (NL4-3ΔCT) by introducing two stop codons at the N’ terminus of the gp41 cytoplasmic tail. We co-transfected 293T cells with either NL4-3 with intact gp41 cytoplasmic tail (HIV-1) or NL4-3ΔCT (HIV-1ΔCT) and either human MARCH1, 2, 8 or E.V. Unlike our findings with MLV, we noticed that the gp41 cytoplasmic tail is dispensable for MARCH-mediated restriction (Fig. 5B). In conclusion, the difference in the importance of the cytoplasmic tail between gp41 and p15E for MARCH-mediated restriction suggests that MARCH proteins may use different mechanisms to restrict MLV and HIV-1.

### Mouse MARCH1 and 8 physically interact with the p15E subunit of the MLV envelope utilizing different transmembrane domains

MARCH proteins physically interact with their cellular targets (MHC II, IL1RAcP etc.) (24, 39). To determine if endogenous mouse MARCH1 and mouse MARCH8 directly interacted with MLV p15E, we infected MutuDC1940 cells, which express endogenous levels of mouse MARCH1 and mouse MARCH8 (Fig. 1C), with MLV. Cells were lysed and co-immunoprecipitations (coIPs) were performed by immunoprecipitating for either mouse MARCH1 or mouse MARCH8. A p15E specific antibody revealed that MLV p15E co-immunoprecipitated with endogenous mouse MARCH1 (Fig. 6A) and MARCH8 (Fig. 6B). To determine the importance of the TM domains of mouse MARCH1 for binding to MLV p15E, we transfected 293T cells with mouse MARCH1, mouse MARCH1 with the TM domains swapped with those from MARCH3, as mouse MARCH3 does not bind to p15E (Fig. S4B), and an MLV molecular clone. We performed coIPs with anti-myc (MARCH1) or anti-p15E. We found that MARCH1 co-immunoprecipitated with MLV p15E independent of the TM domains present (Fig. 6C). MLV p15E also co-immunoprecipitated with wild type MARCH1 as well as MARCH1 containing the MARCH3 TM domains (Fig. 6C). Thus, we concluded that mouse MARCH1 interacts with p15E independently of the TM domains. To determine the role of the TM domains of mouse MARCH8 on its interaction with MLV p15E, we used the mouse MARCH8 constructs with the mouse MARCH4 TM domains mentioned above (see Figure 4F). MARCH4 does not bind to p15E (Fig. S4C). We found that changing the C’ terminal TM domain (TM2) of mouse MARCH8 to that of mouse MARCH4 abolished the mouse MARCH8-p15E interaction (Fig. 6D). In summary, our findings show that mouse MARCH1 and mouse MARCH8 physically interact with the MLV p15E and while for mouse MARCH1, its interaction with p15E is TM-independent, mouse MARCH8 interacts with p15E via the C’ terminal TM domain (TM2).

**Fig 6.**
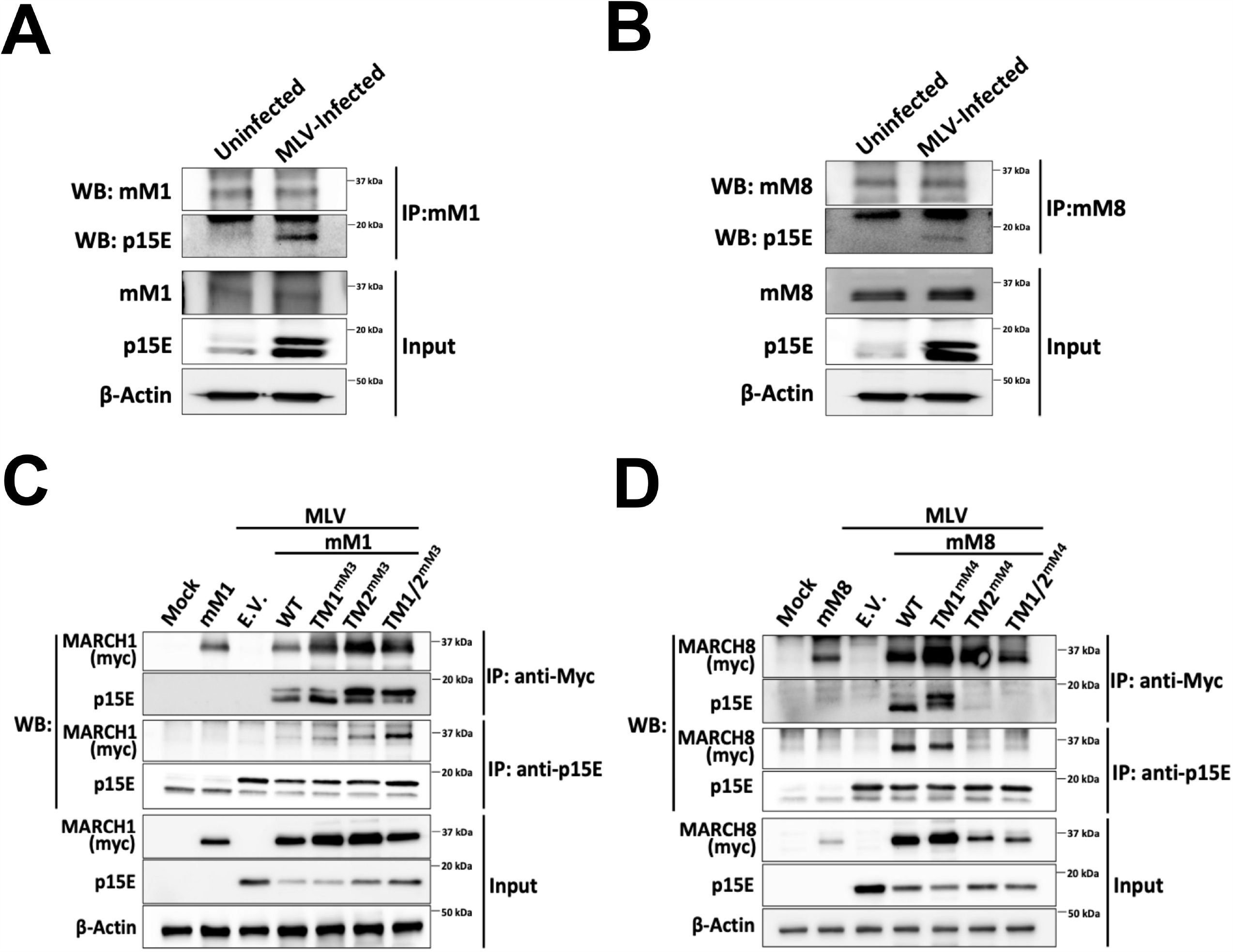
Mouse MARCH1 and mouse MARCH8 physically interact with MLV p15E. (A-B) MutuDC1940 cells, which express endogenous levels of both mouse MARCH (mM) 1 and mM8, were infected with MLV (10 MOI). Cells were harvested 72 hours post infection and lysates were immunoprecipitated with (A) anti-MARCH1 and (B) anti-MARCH8 and analyzed in western blots with anti-p15E, anti-MARCH1, anti-MARCH8 and anti-β-actin. For (C) and (D), 293T were co-transfected with an MLV infectious clone and either for (C) mouse MARCH1 (mM1) or for (D) mM8 and their TM mutants used in Figure 4. Cells were harvested 48 hours post transfection and lysates were immunoprecipitated with anti-myc (mM1 and mM8) and anti-p15E followed by western blots probing with anti-p15E, anti-myc (mM1 and mM8) and anti-β-actin antibodies. Results are shown for n=3 independent experiments. Representative immunoblotting results are shown for A through D.

### Mouse MARCH1 and 8 target the MLV envelope glycoprotein for lysosomal degradation

Targets of MARCH proteins are degraded either in the lysosome or the proteasome (4, 6, 23, 24). To determine where the MLV envelope glycoproteins are degraded by mouse MARCH1 and MARCH8, we used chloroquine and MG132, a lysosomal and a proteasomal inhibitor respectively. 293T cells were co-transfected with either mouse MARCH1 or mouse MARCH8 along and with an MLV expressing plasmid. Six hours post transfection, media was changed and cells were treated with either 100μM of chloroquine or 16μM of MG132. We observed that chloroquine treatment rescued p15E from MARCH-mediated degradation (Fig. 7A). On the other hand, MG132 had no effect on MARCH-mediated degradation of p15E (Fig. 7B). Therefore, we concluded that both mouse MARCH1 and 8 result in the lysosomal degradation of MLV p15E.

**Fig 7.**
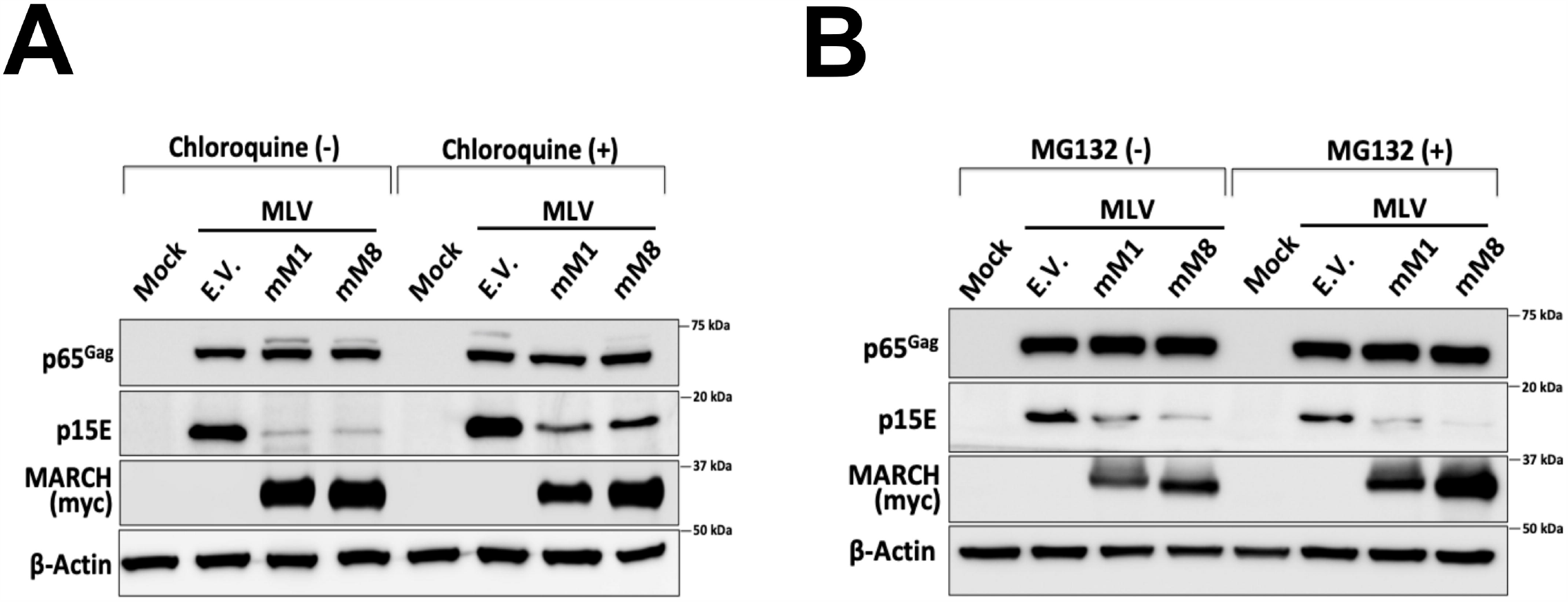
MARCH-mediated degradation of the MLV envelope glycoproteins occurs in the lysosome. (A-B) 293T cells were co-transfected with an MLV infectious clone and either mouse MARCH1 (mM1), mM8 or empty vector (E.V.). Cells were incubated with either (A) 100 μM chloroquine or (B) 16 μM MG132. Cells were harvested 24 hours post transfection and lysates were analyzed by immunoblotting using anti-MLV p30 (detects p65^Gag^), anti-MLV p15E, anti-myc (for mM1 and 8 detection) and anti-β-actin antibodies. Results are shown for n=3 independent experiments. Representative immunoblotting results are shown for A and B.

### Endogenous mouse MARCH1 and MARCH8 restrict retrovirus infection

To determine the role of endogenous mouse MARCH1 and 8 on retrovirus restriction, we transfected BMDCs, which express endogenous mouse MARCH1 and 8 (Fig. 1A and B), with a mouse MARCH1 or 8 specific siRNA and 40 hours post transfection infected them with MLV (0.1 MOI). DNA, RNA and protein lysates of the infected cells were analyzed by RT-qPCR (Fig. 8 A, B and C) and western blots (Fig. 8D) at 6 and 24 hours post infection to measure MLV DNA levels and to ensure efficient MARCH1 and 8 knockdown. In the case of MARCH1, we found that at 24 hours post infection the MARCH1 knockdown infected BMDCs had twice as much MLV DNA compared to those transfected with a control siRNA (Fig. 8A). For MARCH8, we noticed that at 6 hours post infection, MARCH8 knockdown BMDCs had significantly higher levels (∼10X) of MLV DNA than BMDCs transfected with a control siRNA (Fig. 8B). We attempted to look at later time points with the infected MARCH8 knockdown BMDCs, but we noticed significant levels of cell death and only a small number of infected cells survived at 24 hours post infection (Fig. 8B). It is possible that the presumably high levels of MLV envelope glycoproteins in the MARCH8 knockdown BMDCs may be toxic to the cells. Finally, we also verified mouse MARCH1 and 8 were efficiently knocked down (Fig. 8C, D). We concluded that while both endogenous mouse MARCH1 and 8 restrict MLV infection, endogenous mouse MARCH8 is more efficient in inhibiting MLV infection compared to mouse MARCH1.

**Fig 8.**
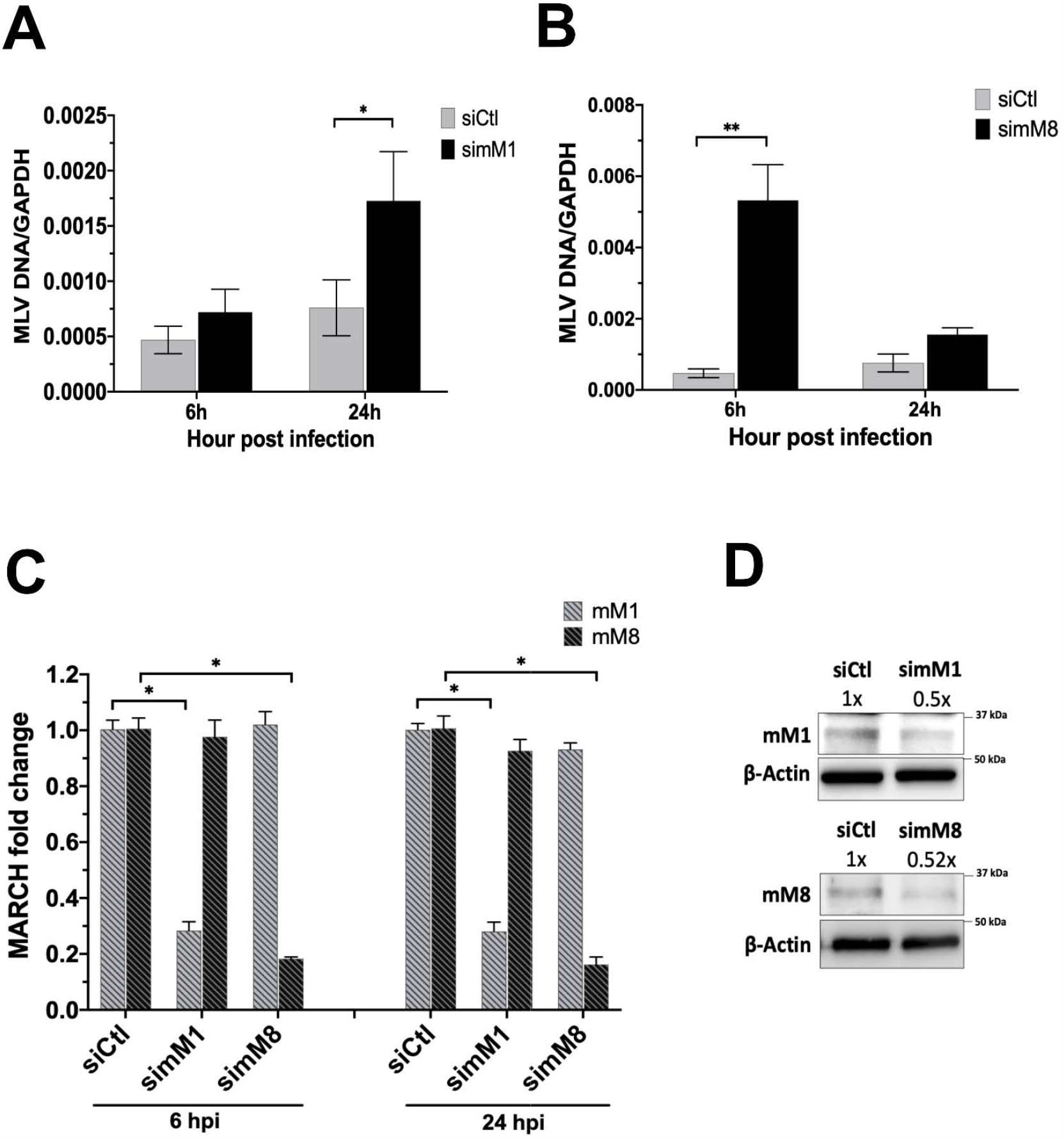
Endogenous mouse MARCH1 and mouse MARCH8 restrict MLV infection. (A-B) Bone marrow derived dendritic cells (BMDCs) were transfected with the indicated siRNAs and infected with MLV (0.1 MOI). Cells were harvested 6 and 24 hours post infection and MLV DNA levels were examined by RT-qPCR. (C-D) Knockdown verification of mouse MARCH1 (mM1) and mM8 from siRNA transfected BMDCs relative to siControl (siCtl) in (A) and (B) by either (C) RT-PCR or (D) immunoblotting. Data shown represent averages of results from n=5 independent experiments. Results for (A) through (C) are presented as means ± standard error of the mean (SEM). Representative immunoblotting results are shown for D, ImageJ (NIH) was used for quantitation of the mM1 and mM8 protein levels. Statistical analysis performed using one sample t-test and Wilcoxon signed rank test and unpaired t-test. * = p ≤ 0.05, ** = p ≤ 0.01.

### MARCH proteins potently restrict a large number of disparate viral envelope glycoproteins

We also investigated whether MARCH proteins have a broad antiviral effect. To address this, we co-transfected 293T cells with either human MARCH1, 2, or 8 and various constructs expressing a number of different viral envelope glycoproteins. Transfected cells were subsequently lysed and western blots were performed to determine viral envelope glycoprotein levels. We first examined the effect of human MARCH1, 2 and 8 on Ebolavirus (EBOV), a member of the filovirus family. We observed that the levels of the mature GP_2_ subunit of the EBOV envelope glycoprotein were reduced in the presence of human MARCH1, 2 and 8 (Fig. 9A). Similar results were seen with LCMV, LASV, and Junín virus JUNV, three member of the arenavirus family, where human MARCH1, 2 and 8 potently reduced the levels of LCMV (Fig. 9B), LASV (Fig. S5A) and JUNV GP_2_ (Fig. S5B), a subunit of the mature envelope glycoprotein complex of arenaviruses (40, 41). We also examined the effect of human MARCH1, 2 and 8 on Nipah Virus (NiV), a highly pathogenic paramyxovirus. We found that human MARCH1, 2 or 8 reduced the levels of NiV fusion (F) and attachment (G) envelope glycoproteins (42) in the transfected cells (Fig. 9C). Additionally, we investigated the effect of human MARCH1, 2 and 8 on hemagglutinin (HA), a viral envelope glycoprotein of orthomyxoviruses, of the lab prototype Influenza A virus (IAV) strain A/PR/8/34 (H1N1). We noticed that human MARCH1, 2 and 8, potently reduced the protein levels of IAV HA (Fig. 9D). Finally, in the case of the envelope glycoproteins E1 and E2 of Chikungunya virus (CHIKV), an alphavirus and a member of the togavirus family, only human MARCH2 and MARCH8 reduced the protein levels of both E1 and E2 (Fig. 9E).

**Fig 9.**
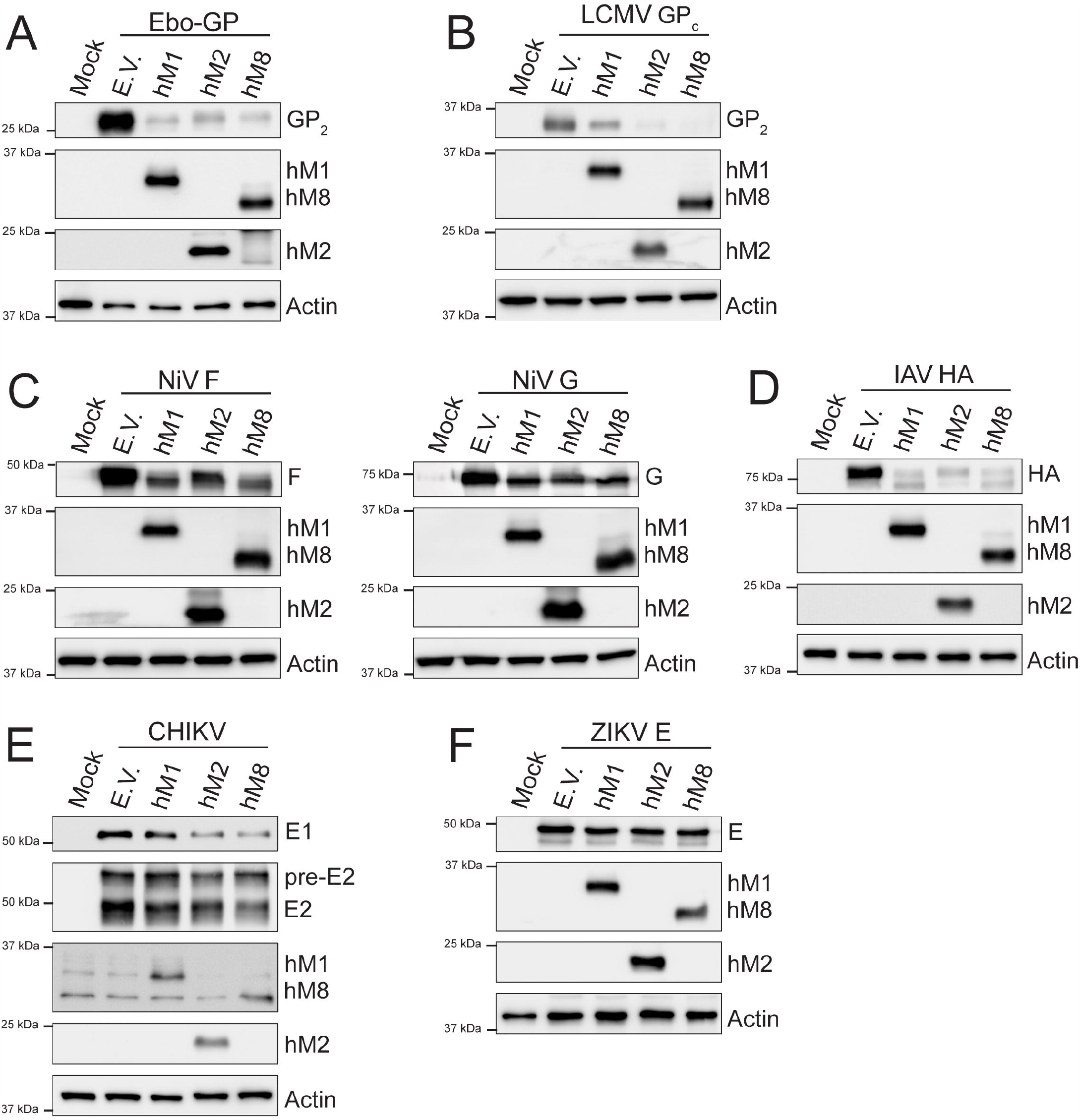
MARCH proteins target envelope glycoproteins from a diverse number of viral families. (A) Ebola virus (EBOV) mature envelope glycoprotein Gp_2_, (B) Lymphocytic Choriomeningitis virus (LCMV) mature envelope glycoprotein GP_2_, (C) Nipah Virus (NiV) fusion glycoprotein (F) (left) and NiV receptor binding glycoprotein (G) (right), (D) Influenza A Virus (IAV) hemagglutinin (HA), (E) Zikka virus (ZIKV) E and (F) Chikungunya virus (CHIKV) E1 and E2 protein levels in the presence of human MARCH1 (hM1), MARCH2 (hM2), MARCH8 (hM8) and empty vector (E.V). 293T cells were co-transfected with either (A) EBOV glycoprotein (Gp), (B) LCMV Gp, (C) NiV F or NiVG, (D) IAV HA, (E) a plasmid expressing the CHIKV structural proteins or (F) ZIKV E in combination with either hM1, 2, 8 or E.V. At 24 hours post transfection, cells were lysed and analyzed by western blots using anti-V5 (for EBOV GP_2_ and IAV HA detection), anti-FLAG (for LCMV GP_2_ detection), anti-AU1 (for NiV G detection), anti-HA (for NiV F detection), anti-Flavivirus E antibody (4G2-for ZIKV E detection), anti-myc (CHIKV E1 detection), anti-E2 (CHIKV E2 detection), anti-MARCH1, anti-MARCH2, anti-MARCH8 and anti-β-actin antibodies. Shown are the results of a single experiment (representative of three independent experiments).

While we observed a fairly broad antiviral effect by human MARCH1, 2 and 8 against a variety of viral envelope glycoproteins, we also identified a number of viral envelope glycoproteins that were resistant to human MARCH1, 2 and 8-mediated degradation. In the case of two members of the paramyxovirus family we studied, we found that both Human Parainfluenza virus 1 (HPIV-1) Hemagglutinin-neuraminidase (HN) and Measles virus (MV) hemagglutinin (H) were resistant to human MARCH1, 2 and 8-mediated degradation (Fig. S5C and S5D). Moreover, the envelope glycoproteins of Zika virus (ZIKV) and West Nile virus (WNV), two members of the flavivirus family, were also resistant to MARCH1, 2 and 8-mediated degradation (Fig. 9F and S5E). Finally, the Gc envelope glycoprotein of Crimean-Congo Hemorrhagic Fever virus (CCHFV), a member of the orthonairovirus family, was also unaffected by human MARCH1, 2 and 8 (Fig. S5F). In summary, our findings above show that MARCH proteins can restrict viral envelope glycoproteins of a number of viruses outside of the retrovirus family, further emphasizing their role as important antiviral factors.

A novel pathogen was recently discovered that was subsequently named Severe Acute Respiratory Syndrome Coronavirus 2 (SARS-CoV-2) (43, 44). Consequently, we examined the effect of human MARCH1, 2 and 8 on the SARS-CoV-2 viral envelope glycoproteins-the S protein, which is important for receptor binding and fusion (45), and the M protein found at high levels on the viral envelope (46), which shares 90% sequence similarity with the M protein of SARS CoV (47), is O’ and N’ glycosylated and drives coronavirus assembly (48, 49). We found that human MARCH2 and 8 reduced S protein levels in the transfected cells, while MARCH1 had no effect (Fig. 10A). Similar to previous reports, we observed 3 forms of the SARS-CoV-2 M protein (50). A prominent diffuse band at 30-50KDa, which likely represents a heterogeneous population of glycosylated SARS-CoV-2 M that is incorporated in the nascent virions and aids in immune evasion, and two distinct forms of M protein that migrate with an apparent molecular mass of 18 and 22KDa respectively (Fig. 10B) (50). Our data show that human MARCH1, 2 and 8 targeted specifically the highly glycosylated forms of SARS CoV-2 M that migrate at 30-50KDa, while the lower molecular weight forms of SARS-CoV-2 M remained unaffected (Fig. 10B). In conclusion, we show that human MARCH2 and 8 cause the degradation of two SARS-CoV-2 envelope glycoproteins, M and S, whereas MARCH1 only targets SARS-CoV-2 M for degradation.

**Fig 10.**
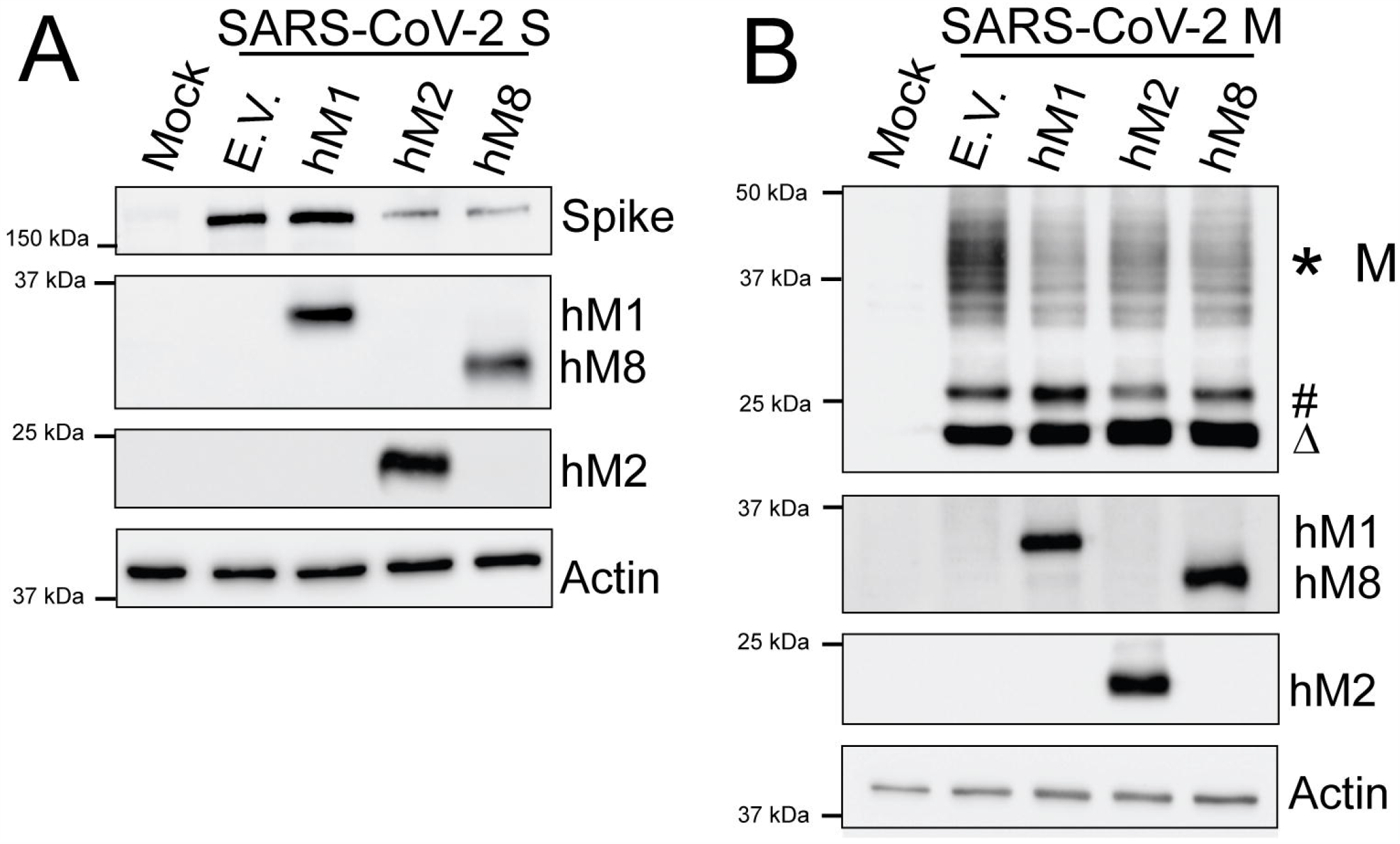
MARCH proteins potently restrict the SARS-CoV-2 Spike (S) and Membrane (M) envelope glycoproteins. (A) SARS-CoV-2 Spike (S) and (B) SARS-CoV-2 Membrane (M) cellular levels in the presence of human MARCH1 (hM1), MARCH2 (hM2), MARCH8 (hM8) and empty vector (E.V.). 293T cells were co-transfected with SARS-CoV-2 S (A) or M (B) and either hM1, 2, 8 or E.V. At 24 hours post transfection, cells were lysed and analyzed by western blots using anti-SARS-CoV-2 S, anti-V5 (for SARS-CoV-2 M detection), anti-MARCH1, anti-MARCH2, anti-MARCH8 and anti-β-actin antibodies. The *, # and Δ symbols represent different glycosylated forms of SARS-CoV-2 M. Shown are the results of a single experiment (representative of three independent experiments).

## Discussion

MARCH proteins are membrane associated E3 ubiquitin ligases, whose main function is to tightly regulate the levels of immune receptors by targeting them for degradation (e.g. MHC II, Transferrin receptor, CD86 etc.) on the PM (1, 4, 9, 19, 51, 52). Human MARCH1, 2 and 8 also target the HIV-1 envelope glycoproteins at the PM leading to their intracellular sequestration blocking their incorporation in nascent virions (30-32). Nevertheless, many questions remain regarding the antiviral mechanism of these host factors.

We initially examined the effect of IFN-β and virus infection on mouse MARCH1, 2, 3 and 8 and found that only mouse MARCH1 was induced by IFN-β, while MLV infection had no effect on mouse MARCH1, 2, 3 and 8 expression. Furthermore, we found that only mouse MARCH1 and 8 caused the degradation of the viral envelope glycoproteins of MLV and MMTV. Our findings suggest that MARCH proteins utilize a similar mechanism to target viral envelope glycoproteins and immune receptors for removal from the PM followed by degradation. Therefore, the ability of MARCH proteins to remove and degrade proteins from the cell surface is utilized by the cell in two ways: (1) for the maintenance of homeostasis, by regulating receptor levels in the PM, and (2) for the removal and degradation of viral envelope glycoproteins as means to thwart viral infection.

As we noticed the degradation and not the internalization and sequestration of the retroviral envelope glycoproteins, as previous studies showed, we compared side by side the effect of human and mouse MARCH1, 2 and 8 against both MLV and HIV-1 envelope glycoproteins. One of the key observations we made was that mouse MARCH2, unlike human MARCH2, had no antiviral effect. This is particularly interesting as mouse MARCH2 and human MARCH2 share more than 80% sequence homology (data not shown). The reason for the discrepancies in both antiviral function and transcriptional regulation between the human and mouse homologues of MARCH2 is currently unknown. Furthermore, MARCH1 also displayed a very interesting antiviral phenotype; mouse MARCH1 was only antiviral against MLV and MMTV but not against HIV-1, whereas human MARCH1 potently downregulated envelope glycoproteins of all retroviruses tested. This suggests that human MARCH1 has acquired broader antiviral functions at a later point in the evolutionary scale allowing it to target a greater variety of retroviral envelope glycoproteins (HIV-1 and MLV). The reason behind the differences in restrictive range between mouse and human MARCH1 needs to be further elucidated. In the case of MARCH8, both mouse and human homologues degraded all retroviral envelope glycoproteins tested. To summarize, our findings suggest that MARCH proteins are important antiviral factors that are under constant selective pressure to counteract retrovirus infections.

Mouse MARCH1 and MARCH8 are multipass TM proteins that have both their N’ and C’ termini inside the cytoplasm (1, 15). Many domains in MARCH1 and MARCH8 have been previously characterized as essential for their ability to remove proteins from the PM. In this report, we identified that only the RING-CH and DIRT domains found in the N’ terminal cytosolic tail are essential for restricting viral envelope glycoproteins. Moreover, while the C’ terminal TM domain is important for mouse MARCH1 restriction, it did not affect the MARCH1-MLV p15E interaction. On the other hand, the C’ terminal TM domain of mouse MARCH8 is critical for restriction and interaction with MLV p15E. Our data thus suggest that mouse MARCH1 and 8 may utilize different mechanisms to restrict MLV p15E.

We also show that the cytoplasmic tail of the MLV p15E, unlike that of HIV-1, is critical for MARCH-mediated removal from the PM, similar to cellular immune receptors targeted by MARCH proteins (6, 18, 19, 25, 53). The variability in the importance of the retroviral cytoplasmic tail on MARCH-mediated restriction further suggests that MARCH proteins may utilize distinct mechanisms to restrict envelope glycoproteins from different retroviruses.

We also examined the role of MARCH1, 2 and 8 on the viral envelope glycoproteins from a diverse number of enveloped viruses. We found that all three target the viral envelope glycoproteins of a diverse number of viruses, including NiV, EBOV, LCMV and SARS-CoV-2. On the other hand, a number of viral envelope glycoproteins, including that of ZIKV and WNV were unaffected. However, it does not necessarily mean that MARCH proteins do not restrict them; it is possible that they are sequestered intracellularly, blocking their incorporation in nascent virions or that other members of the MARCH family are important in the degradation of these viral glycoproteins. In the case of SARS-CoV-2, we observed that human MARCH2 and 8 target both the S and M proteins while human MARCH1 targets solely the M protein for degradation. The fact that two different envelope glycoproteins of SARS-CoV-2 are restricted by a different gamut of MARCH proteins provides support to the idea that different MARCH proteins may have variable range of targets, something that needs to be further examined. Finally, in the case of SARS-CoV-2 M we see that MARCH proteins only target the heavily glycosylated form, which is the one that is incorporated into virions. This may be due to the fact that the heavily glycosylated form of SARS-CoV-2 M may co-localize with MARCH1, 2 and 8 at the Golgi complex (1, 9, 54) while the non-glycosylated SARS-CoV-2 M forms do not. In summary, we show that the antiviral function of MARCH proteins is well conserved among different species and is quite expansive.

## Materials and Methods

### Cell lines

293T cells (ATCC) and 293FT cells (Invitrogen) were cultured in Dulbecco’s modified Eagle media (DMEM; Gibco) with 10% (vol/vol) fetal bovine serum (FBS; Sigma), 0.1 mM nonessential amino acids (Gibco), 6 mM L-glutamine (Gibco), 100 mg/ml penicillin and streptomycin (P/S; Gibco), 1 mM sodium pyruvate (Gibco), and 500 μg/ml Geneticin (Gibco). NIH 3T3 and *Mus dunni* cells (ATCC) were cultured in DMEM with 10% FBS and P/S. EL4 cells (ATCC) were cultured in RPMI media with 10% FBS, P/S, and 0.05 mM β-mercaptoethanol (β-ME; Bio-Rad). MutuDC1940 cells (33) were cultured in Iscove’s modified Dulbecco’s media (IMDM) with 8% FBS, 100 mg/ml P/S, 1 mM sodium pyruvate, 10 mM HEPES (Corning), and 0.05 mM βME. BMDCs and BMDMs were generated from 6-9 week C57BL6/N mice as previously described (55). BMDCs were differentiated with recombinant murine granulocyte-macrophage colony-stimulating factor (20 ng/ml; Invitrogen), while BMDMs with macrophage colony-stimulating factor (10ng/ml; Invitrogen).

### Plasmids

The MLV molecular clone (pLRB302) and HIV-1 molecular clone (pNL4-3) used in this manuscript have been previously described (56, 57). The pNL4-3 was obtained from NIH AIDS Reagent program, Division of AIDS, NIAID, NIH from Dr. Malcolm Martin (cat#114). To generate an HIV-1 infectious clone lacking the gp41 cytosolic tail (HIV-1ΔCT), we initially digested the pNL4-3 plasmid with *Xho*I and *Eco*RI. We introduced in the NL4-3*Xho*I-*Eco*RI fragment two stop codons at residues 705 and 707 (R705Stop and R707Stop) of the envelope ORF located at the N’ terminus of the cytosolic tail, using the Phusion Site-Directed Mutagenesis Kit (Thermo Fisher Scientific). The primers used to introduce the changes (underlined) were as follows: 5’-CTTTCTATAGTGAAT<u>T</u>GAGTT<u>TA</u>GCAGGGATATTCAC-3’/ 5’-TACAGCAAAAACTATTCTTAAACCTACCAAGC-3’. After sequence verification, the *Xho*I-*Eco*RI fragment was reintroduced to the pNL4-3 backbone. To generate an MLV infectious clone with the cytosolic tail deleted (MLVΔCT), we introduced a stop codon in the pLRB302 plasmid at residue 640 of the envelope ORF (C640Stop) located at the N’ terminal of the cytosolic tail using the Phusion Site-Directed Mutagenesis Kit (Thermo Fisher Scientific) and the following primers: 5’-CTTTTTGGACCCTG<u>A</u>ATTCTTAATCGATTAGTTC-3’/ 5’-CAGAATTAGTAGGAGTATAATGAGAGGCCCCAT-3’. (The change introduced is underlined in the forward primer). The presence of the desired mutations was verified by sequencing.

Mouse MARCH1, 2, 3, 4 and 8 cDNA clones were purchased from Dharmacon (MMM1013-202858906, MMM1013-202761409, MMM1013-211693113, MMM1013-202799076 and MMM1013-202798072 respectively). Mouse MARCH1, 2, 3, 4 and 8 were initially cloned into pCDNA3.1/myc-His A (Invitrogen) and were later subcloned into pBJ5 vector using the NEBuilder HiFi DNA assembly kit (New England Biolabs). To amplify mM1, 2, 3, 4 and 8 genes from pcDNA3.1/myc-His A, the same reverse primer was used for all mouse MARCH genes: 5’-GGCCTCCGCGGCCGCTCAATGGTGATGGTGATGAT- 3’, while the forward primers were different for all genes and are the following: for mM1, 5’-GCTCTAGCCCTCGAGATGCCCCTCCACCAGATTTC- 3’/ for mM2, 5’-GCTCTAGCCCTCGAGATGACGACAGGTGACTGTTGCC- 3’/ for mM3, 5’-GCTCTAGCCCTCGAGATGACAACCAGTCGCTGCAGTC- 3’/ for mM4, 5’-GCTCTAGCCCTCGAGATGCTCATGCCCCTGGGTGG- 3’/ for mM8, 5’ GCTCTAGCCCTCGAGATGAGCATGCCATTGCACCAG-3’. To amplify the pBJ5 vector, the following forward primer was used for all mouse MARCH genes: 5’- ATCACCATTGAGCGGCCGCGGAGGCCGAATTC-3’. The reverse primers used to amplify pBJ5 differed among the different mouse MARCH genes and are the following: for mM1, 5’- TGGTGGAGGGGCATCTCGAGGGCTAGAGCAGCTTTTAGAG-3’/ for mM2, 5’-GGCAACAGTCACCTGTCGTCATCTCGAGGGCTAGAGCAGCTTTTAGAG- 3’/ for mM3, 5’-GACTGCAGCGACTGGTTGTCATCTCGAGGGCTAGAGCAGCTTTTAGAG- 3’/ for mM4, 5’-CCACCCAGGGGCATGAGCATCTCGAGGGCTAGAGCAGCTTTTAGAG- 3’/ for mM8, 5’ CTGGTGCAATGGCATGCTCATCTCGAGGGCTAGAGCAGCTTTTAGAG-3’. PCR products were amplified and recombined using the NEBuilder HiFi DNA assembly kit (New England Biolabs) followed by Sanger sequencing for identification of positive clones. In the case of human MARCH genes, hM1 cDNA was synthesized from HeLa cell RNA. The pCMV-hM2-3HA has been previously described (31) and kindly provided to us by Xinqi Liu, while cDNA of hM8 was obtained from Dharmacon (MHS6278-202807570). All human MARCH cDNAs were cloned into pCDNA3.1/myc-His A (Invitrogen) and were then subcloned into pBJ5. hM1, 2 and 8 were initially PCR amplified using a common reverse primer: 5’-GGCCTCCGCGGCCGCTCAATGGTGATGGTGATGAT- 3’ and the following forward primers: for hM1, 5’-GGCTCGAGATGCTGGGCTGGTGTGAAGCG -3’/ for hM2, 5’- GGCTCGAGATGACGACGGGTGAC -3’/ and for hM8, 5’-GGCTCGAGATGAGCATGCCACTG -3’. The PCR fragments and pBJ5 were cut with *Not*I and *Xho*I followed by ligation, transformation and sequencing for positive clones. To generate all the mM1 and mM8 mutants described in this manuscript, we used the Phusion Site-Directed Mutagenesis kit (Thermo Fisher Scientific). To generate the RING-CH^mut^ mM1 and mM8, mutations were introduced at C80S, C83S, C97S and C99S using the primers described in Table I. To generate mM1 and mM8 ΔDIRT, we deleted residues 130 to 140 using the primers described in Table I. To mutate the tyrosine endocytic motifs (YXXΦ) of mM1 (^218^YVQL^221^and ^228^YNRL^231^) and mM8 (^218^YLQL^221^ and ^228^YNRL^231^), we mutated them to ^218^AAQL^221^ and ^228^AARL^231^ for both mM1 and mM8 using the primers shown in Table I. The ^234^VQNC^237^ motifs of mM1 and mM8 were mutated to ^234^AANC^237^ using the primers mentioned in Table I. To generate mM1 and mM8 carrying the TM domains of mM3 and mM4 respectively, we initially PCR amplified the TM domains of mM3 and mM4 with the primers shown in Table I. We then PCR amplified mM1 and mM8 with primers flanking their TM domains and described in Table I. PCR products recombined using the NEBuilder HiFi DNA assembly kit (New England Biolabs). All constructs were verified by sequencing.

**Table I.**
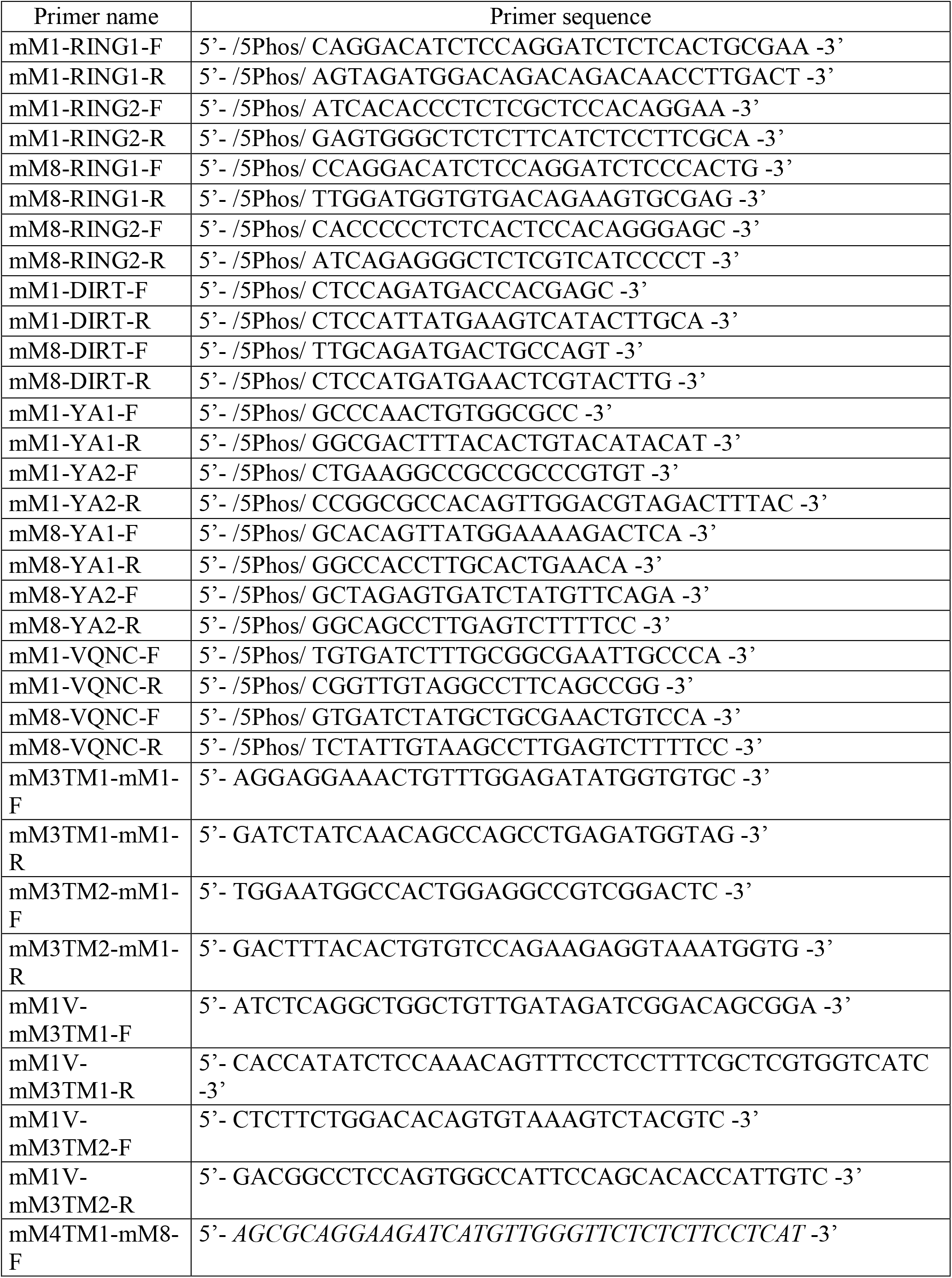
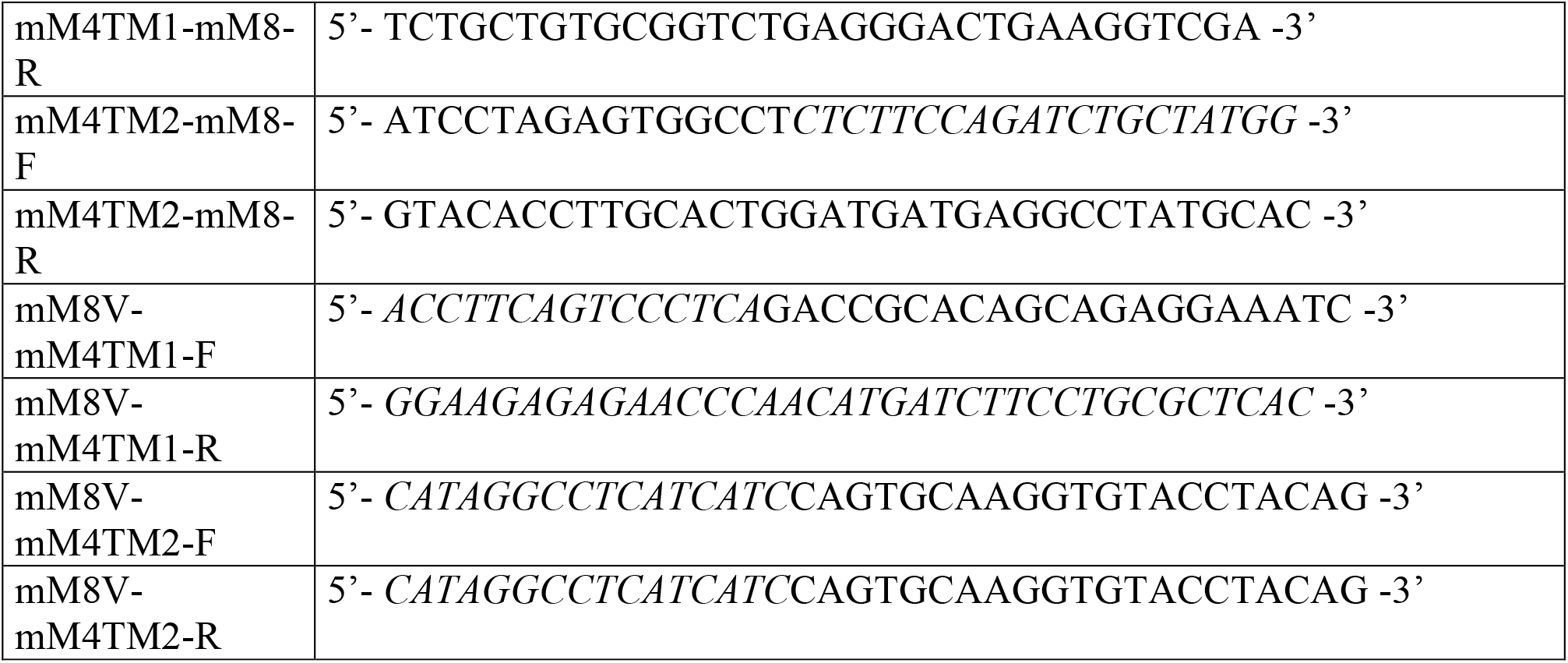
Primers for making mM1 and mM8 mutants.

Influenza A virus (IAV) hemagglutinin (HA) gene was PCR amplified from cDNA generated from IAV A/Puerto Rico/8/1934 (H1N1) genomic RNA (BEI Resources, NIAID, NIH, NR-2773) using the primers described in Table II followed by restriction digestion with *Bam*HI and *Xba*I and ligation into pcDNA-V5/His (Invitrogen). The MV H and HPIV-1 HN were PCR amplified using the primers shown in Table II from genomic RNA of MV, Edmonston strain and from virus of the HPIA1-HN-F/HPIA1-HN-R strain of HPIV-1 (BEI Resources, NIAID, NIH, NR-44104 and NR-48681 respectively), and cloned into pcDNA-V5/His (Invitrogen). The M fragment of CCHFV was PCR amplified with primers shown in Table II from cDNA generated from CCHFV genomic RNA, IbAr10200 strain (BEI Resources, NIAID, NIH NR37382). PCR products were cloned into pcDNA-V5/His (Invitrogen) using the NEBuilder HiFi DNA Assembly Master Mix (New England Biolabs). SARS-CoV-2 M was PCR amplified from cDNA derived from genomic RNA of SARS-CoV-2, Isolate USA-WA1/2020 (BEI Resources, NIAID, NIH, NR-52285) using the primers in Table II and cloned into pCDNA-V5/His TOPO (Invitrogen). We introduced a FLAG tag to JUNV GP by PCR amplifying it from a previously described expression plasmid (58, 59) using the primers in Table II and after digestion with *Bam*HI and *Xho*I was cloned into pCDNA™3.1/myc-His A (Invitrogen). CHIKV structural genes were initially PCR amplified from pDONR21-CHKVstr (60) kindly provided to us by Ted Pierson, using the primers shown in Table II and cloned into pcDNA-V5/His (Invitrogen). pCAGGS-EboGP-V5 (strain Zaire 1976 Mayinga) (61) and pcDNA V5/His SARS-Cov2 Spike (Wuhan-Hu-1) were kindly provided by Paul Bates. LASV and LCMV GP tagged with a FLAG tag have been previously described (62, 63). pcDNA 3.1(+) ZIKA ENV was provided by Amy Jacobs. Plasmids encoding codon optimized NiV F and NiV G genes tagged with AU1 and HA tag respectively were kindly provided by Benhur Lee (64). pVAC2-WNV prME plasmid was kindly given to us by Ted Pierson (65).

**Table II.**
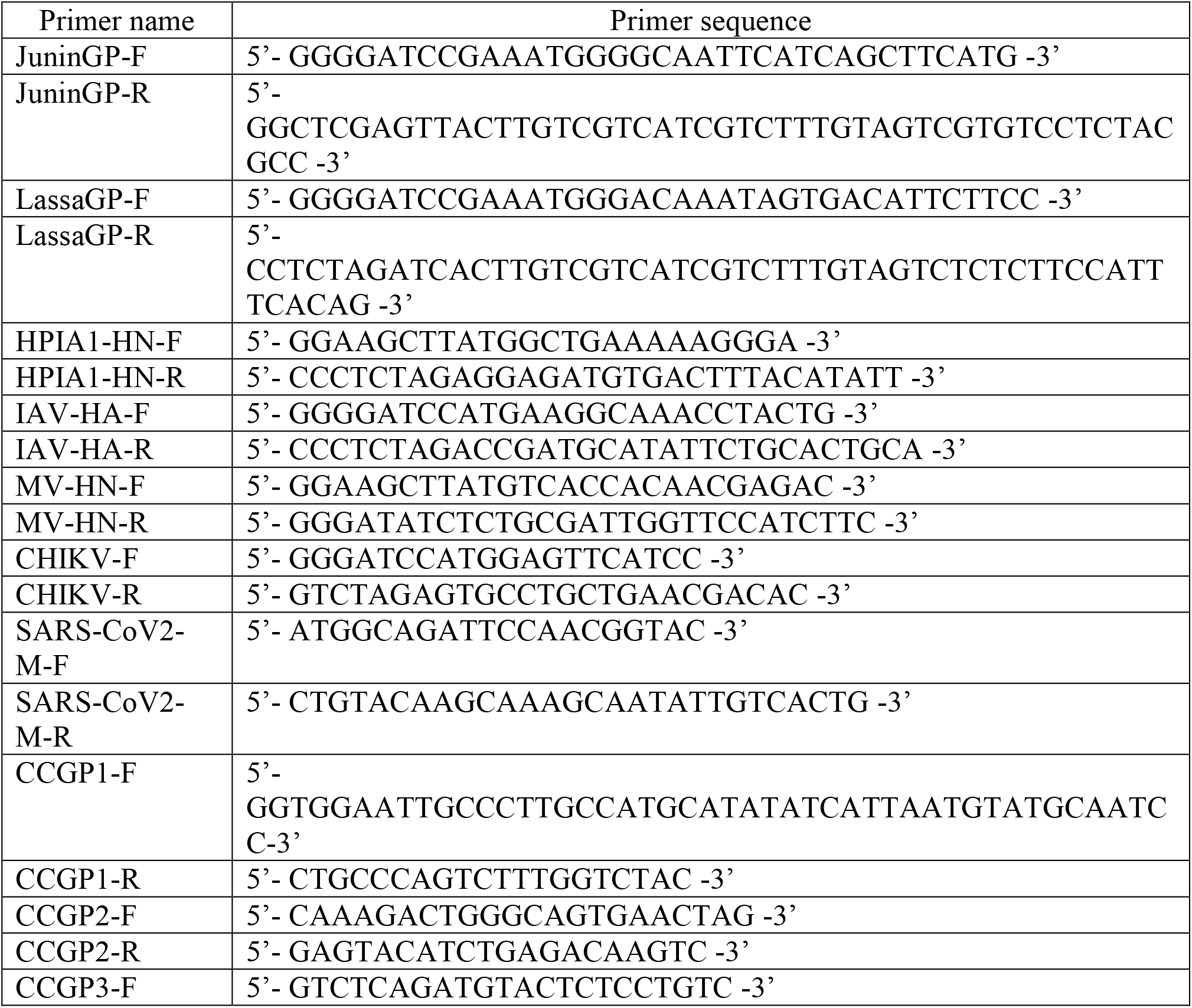
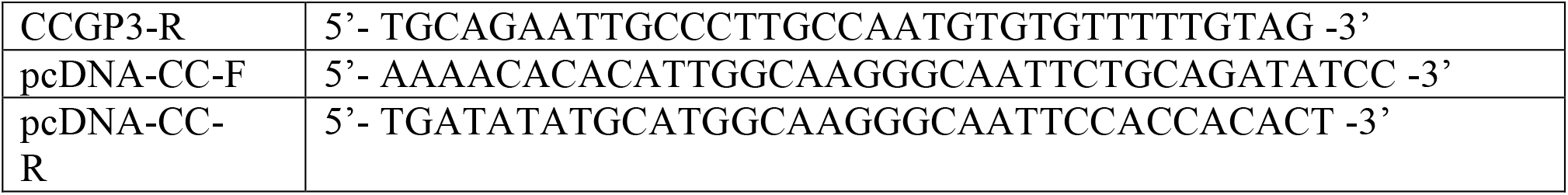
Primers for cloning of viral envelopes.

### Transfections and western blots

For all transfection experiments we used Lipofectamine 3000 transfection reagent (Thermo Fisher Scientific) according to the manufacturer’s recommendation. 293T cells were seeded on a 6-well plate (0.5 × 10^6^cells/well) and the following day were co-transfected with 3 μg of MLV (pLRB3020) or 3 μg of pNL4.3 along with either 4 μg of pBJ5-mMARCHs (mM1, mM2, mM3, or mM8) or with 4 μg of pBJ5-hMARCHs (hM1, hM2, or hM8). To study the dose-dependent effect of mM1 and mM8, we used the following concentrations of mM1 and mM8: 0.25, 0.5, 1, 2, and 4 μg. For the MMTV transfection experiments, we seeded 293T cells in a 6-well plate (0.5 × 10^6^/well) and transfected them with 5 μg of MMTV hybrid provirus (HP) plasmid (34), 50ng of rat glycocorticoid receptor (RSVGR) construct and 1 μg mM1, 2, 3, 8 or E.V (1 μg). For the co-transfection experiments with the mutant mM1 and mM8 constructs, we seeded 0.5×10^6^ 293T cells and the following day we co-transfected them with 3 μg of MLV infectious clone (pLRB302) and 1 μg of either E.V., mM1, mM8, or mM1 and mM8 with mutations in the various domains mentioned in the Plasmids section. To determine the role of the cytosolic tail of MLV p15E and HIV-1 gp41, we co-transfected 1 μg of mM1 or mM8 with either 3 μg of wild type MLV, wild type HIV-1, MLVΔCT or HIV-1ΔCT. For all above transfections, cells and culture media were harvested 48 h after transfection. Cells were lysed in RIPA buffer (150mM NaCl, 1% NP-40, 0.5% Sodium Deoxycholate, 0.1% SDS, 25mM Tris pH7.4 with HALT™ phosphatase and protease inhibitors) and viruses from the culture media were used for future experiments. Cell lysates and virus pellets were mixed with 1× sample loading buffer before they were resolved on 10% or 15% sodium dodecyl sulfate polyacrylamide gels. Blots were probed using the following antibodies: goat anti-MLV gp70 (66), goat anti-MMTV polyclonal (67), rat anti-MLV transmembrane protein/p15E (clone 42/114, Kerafast), rabbit anti-myc (Cell Signaling Technology), rat anti-MLV p30 (R187, ATCC CRL-1912), mouse anti-HIV-1 envelope (16H3 NIH/AIDS Reagent program Cat#12559), mouse anti-HIV gp41 (Chessie8) (NIH/AIDS Reagent program Cat#526), human anti-HIV gp41 clone 2F5 (NIH/AIDS Reagent program Cat#1475) for detection of gp41ΔCT, mouse anti-HIV p24 (NIH/AIDS Reagent program Cat#4121), rabbit anti-MARCH1 (Thermo Fisher Scientific Cat#PA5-20632), rabbit anti-MARCH2 (Thermo Fisher Scientific Cat#PA5-30220), rabbit anti-MARCH8 (Thermo Fisher Scientific#PA5-20632), and monoclonal anti-β-actin (Sigma-Aldrich). HRP-conjugated anti-rabbit IgG (Cell Signaling Technology), HRP-conjugated anti-rat IgG (Cell Signaling Technology), HRP-conjugated anti-human IgG (Sigma Aldrich, Cat#GENA933), HRP-conjugated anti-mouse (EMD Millipore), and HRP-conjugated anti-goat (Sigma-Aldrich) were used for detection using the enhanced chemiluminescence detection kits Clarity and Clarity Max ECL (Biorad).

To determine the effect of MARCH proteins on the non-retroviral envelope glycoproteins, we seeded 8×10^5^ 293T cells in a 6 well plate and co-transfected them with either 4 μg of hM1, 2, 8 or E.V. and 50 ng of EBOV GP-V5, LCMV GP-FLAG, ZIKV E, LASV GP-FLAG, SARS-CoV-2 M-V5, SARS-CoV-2 S. In the case of IAV HA, 4 μg of hM1, 2, 8 or E.V. were co-transfected with 100 ng of IAV HA-V5. For NiV F and G, 25 ng of either NiV F-AU1 or NiV G-HA were co-transfected with 4 μg of either hM1, 2, 8 or E.V. For CCHFV Gc, CHIKV E1 and E2, MV H and HPIV-1 HN, we co-transfected 4 μg of the viral envelopes along with 700 ng of hM1, 2, 8 or E.V. In the case of WNV, we co-transfected 400 ng of WNV PrmE along with 1 μg of hM1, 2, 8 or E.V. For JUNV, we co-transfected 5 μg of JUNV Gp-FLAG and 1 μg of hM1, 2, 3, 8 or E.V. Cells were harvested 48 hours post transfection and lysed in RIPA buffer except for MV H and HPIV HN, which were lysed in DM lysis buffer [0.5% (w/v) n-Decyl-β-D-Maltopyranoside, 20mM Tris-HCl, pH 7.5, 10% (v/v) glycerol, 1X Halt Protease inhibitor cocktail (Thermo Fisher Scientific), Benzonase (25 U/mL)]. Lysates were then resolved on 10% sodium dodecyl sulfate polyacrylamide gels and blots were probed using the following antibodies: rabbit anti-FLAG (Cell Signaling Technology), rabbit anti-V5 (Invitrogen), mouse anti-Flavivirus envelope 4G2 (BEI Resources, NIH, NIAID NR-50327), rabbit anti-AU1 (Novus Biologicals), mouse anti-CHIKV E2 (BEI Resources, NIH, NIAID NR-44002), mouse anti-SARS-CoV-2 S (Genetex Cat#632604) and the rabbit anti-hM1, 2 and 8 described above. HRP-conjugated anti-rabbit and anti-mouse were used for detection using the chemiluminescence detection kit Clarity ECL (Biorad).

### FACS analysis

The purity of the BMDM and BMDC populations used in our experiments was determined by cell surface staining followed by FACS. For BMDMs, we stained 1×10^5^ BMDMs with 1:50 of FITC rat anti-mouse CD11b (clone M1/70 BDPharmingen™ Cat#557396) in PBS with 2% FBS (staining buffer) for 30 minutes at 4°C. Cells were washed with 500 μL of staining buffer twice and re-suspended in 100 μL of staining buffer and processed using a BD LSRFortessaTM flow cytometer. For BMDCs, 1×10^5^ BMDCs were harvested and stained with 1:25 dilution of APC Hamster anti-mouse CD11c (clone HL3 BDPharmingen™ Cat#550261) in staining buffer, washed, and run through a BD LSRFortessaTM flow cytometer.

To verify that the MARCH1 mutants we generated localized to the plasma membrane, we seeded 0.5×10^6^ 293T cells and the following day, cells were transfected with 3 μg of E.V., mM1, or the mutant forms of mM1 using Lipofectamine 3000 (Thermo Fisher Scientific) per the manufacturer’s recommendation. Cells were harvested and 1 × 10^5^ cells were stained with 1:50 of rabbit anti-MARCH1 (Thermo Fisher Scientific Cat#PA5-69223) or 1:50 of rabbit IgG isotype antibody (Thermo Fisher Scientific Cat#02-6102) for 30 minutes at 4°C. Subsequently, cells were washed twice and stained with 1:50 of goat anti-rabbit IgG (H+L) Alexa Fluor 647 (Thermo Fisher Scientific Cat#A21244) for 30 minutes at 4°C followed by two washes and diluted in 100 μL of staining buffer. Samples were processed using a BD LSRFortessaTM flow cytometer. Stained populations from our FACS experiments were further analyzed using Flowjo software version 10.7.1.

### Membrane fractionation of mM8

5×10^5^ 293T cells were seeded and the next day transfected using Lipofectamine 3000 (Thermo Fisher Scientific) with 3 μg of E.V., wild type mM8, or the mutant mM8 constructs we generated. At 24 hours post transfection, cells were collected and membranes were extracted using the Mem-PER™ plus membrane extraction kit (Thermo Fisher Scientific) according to the manufacturer’s guidelines. The purity of the membrane fractions was verified by western blots probing with rabbit anti-GAPDH (Cell Signaling Technology).

### Virus infection of BMDCs

BMDCs were seeded (5 × 10^4^ cells/well) in 96-well plate. The following day, cells were transfected with 3 pmoles of siRNA Control (Ambion Cat#AM4611), simM1, (Ambion ID: s91382, sense: CGUGUGAUCUUUGUGCAGAtt/antisense: UCUGCACAAAGAUCACACGgt), or simM8 (Ambion ID: s90045, sense: ACUCAAGGCUUACAAUAGAtt/antisense: UCUAUUGUAAGCCUUGAGUct) using Lipofectamine RNAiMax (Thermo Fisher Scientific) according to the manufacturer’s recommendation. At 40 hours post transfection, cells were infected with 0.1 MOI of MLV by spinoculation as previously described (68). Cells were harvested and DNA was isolated using the DNeasy blood and tissue kit (Qiagen) at the indicated time points according to the manufacturer’s instructions. RT-qPCR was performed using Power Up™ SYBR™ Green Master mix kit (Applied Biosystems) and the previously described MLV and GAPDH primers (69). A CFX384 Touch real-time PCR detection system (Bio-Rad) was used for all RT-qPCR assays described in this study.

### siRNA knockdown verification

BMDCs were transfected with the indicated siRNAs as described above and RNA was isolated using the RNEasy mini kit (Qiagen). cDNA was generated using SuperScript III first-strand cDNA synthesis kit (Invitrogen). RT-PCR was performed using the Power Up™ SYBR™ Green Master mix kit (Applied Biosystems) and the following primers: mM1, 5’-TCTGCTCTGTCACGTTCCAC -3’/ 5’-CCTCTGCAGTTGGCAGTGTA-3’, mM8, 5’-CTCTCGCACTTCTATCACGCCA -3’/ 5’-AAGTGGAGGCTTCCTGTGCAGT -3’, GAPDH (described in Virus infection of BMDCs section). RT-PCRs were performed as described above.

To detect the knockdown efficacies at the protein level, cells were lysed with RIPA buffer. Cell lysates were resolved on 10% sodium dodecyl sulfate polyacrylamide gels. Blots were probed with the following antibodies: rabbit anti-MARCH1 (Invitrogen Cat#PA5-69223), rabbit anti-MARCH8 (Invitrogen Cat#30220), monoclonal anti-actin (Sigma-Aldrich). HRP-conjugated anti-mouse (EMD millipore) and HRP-conjugated anti-rabbit (Cell Signaling Technology) were used for detection using the chemiluminescence detection kits Clarity and Clarity ECL (Bio-Rad).

### Interferon treatment of cells

0.5 × 10^4^ cells of MutuDC1940, EL-4, NIH3T3, BMDMs and BMDCs were seeded in a 96-well plate for 24 hours. Cells were then treated ± 500 Units/mL of mouse IFN-β (PBL Assay Science) for 4, 8, 16, and 24 hours. For the titration experiment, 0.5×10^4^ MutuDC1940 were treated with IFN-β concentration ranging from 0.005 to 5000 Units/mL and harvested 4 hours post treatment. Cells were lysed and RNA was isolated using an RNeasy mini kit (Qiagen). cDNA synthesis and RT-PCR were performed as described in the siRNA knockdown verification section using the aforementioned mM1, mM8 and GAPDH primers as well as primers for mM2 (5’-TGCCAGCTGTACTCGGAATG-3’/5’-GCTGCATTGCCATCTGACTC-3’) and for mM3 (5’-ATCAGTCGAGCAGAAGCTGAG -3’/ 5’-AGTGTCAGCCTCGTCACATC -3’).

### Virus preparation

MLV stocks were prepared by transfecting 293FT cells (Invitrogen) seeded in 10-cm-diameter cell culture dishes using Lipofectamine3000 (Thermo Fisher Scientific) with 25 μg of an MLV infectious clone (pLRB302) per manufacturer’s recommendation. Culture supernatants were harvested 48 hours after transfection, filtered, and treated with 10 U/ml DNase I (Roche) for 40 min at 37°C. Titers of viruses were determined in *Mus dunni* cells as previously described (69).

### Infection assays

To examine the effect of MLV infection on MARCH gene expression, 0.5 × 10^4^ cells of MutuDC1940, EL4, NIH3T3 were seeded in 96-well plate followed by infection with MLV (5 MOI) via spinoculation as previously described (68). Following spinoculation, cells were given fresh media and harvested at 4, 8, 16, and 24 hours post followed by RNA isolation using an RNeasy mini Kit (Qiagen) per the manufacturer’s recommendations. Primers for mM1, 2, 3, 8 and GAPDH detection are mentioned above.

### Co-Immunoprecipitations

For our co-immunoprecipitation experiments examining endogenous mM1 and 8, we seeded 2 × 10^5^ MutuDC1490 cells in a 6-well plate. The next day, cells were either mock infected (media) or infected by spinoculation with MLV (10 MOI) as mentioned above. Three days post infection, cells were washed once with cold 1X PBS and harvested using NP40 lysis buffer (50mM Tris-HCl, 150 mM NaCl, 1% NP-40, 5mM EDTA, 5% glycerol). Co-Immunoprecipitation was performed using Dynabeads Protein A Immunoprecipitation kit (Thermo Fisher Scientific) following manufacturer protocol with some modifications. Briefly, 50 μl proteinA Dynabeads (Thermo Fisher Scientific) were pre-incubated with 1:50 dilution of rabbit anti-MARCH1 (Invitrogen Cat#PA5-69223) or 1:25 dilution of rabbit anti-MARCH8 (Proteintech Cat#14119-1-AP) antibodies. Cell lysates were incubated with antibodies-coated protein A dynabeads overnight at 4°C, washed, and eluted. The eluted fractions were subjected to SDS-PAGE and western blot analysis. For our co-immunoprecipitation experiments examining the role of the mM1 and 8 TM domains, we co-transfected 293T cells with 3 μg of an MLV infectious clone (pLRB302) and 100 ng of the indicated different mouse MARCH constructs. At 24 hours post transfection cells were lysed in NP40 lysis buffer. Protein A Dynabeads were pre-incubated with anti-Myc (Cell Signaling Technology), or 1:20 of culture supernatant of 372 (ATCC CRL-1893) and then cell lysates were added to antibodies-bound protein A dynabeads and incubated at room temperature for 1 hour, washed, and eluted, followed by SDS-PAGE and western blot analysis detecting for MLV p15E or MARCH(myc).

### Luciferase assays

293T cells were seeded in a 6-well plate (0.5 × 10^6^ cells/well) a day prior to transfections. The following day, co-transfection were performed using 3 μg of an MLV infectious clone (pLRB302), 1 μg of pFB-Luciferase (pFB-luc) plasmid and 4 μg of either pBJ empty vector, pBJ5-mM1, pBJ5-mM2, pBJ5-mM3, or pBJ5-mM8, using Lipofectamine 3000 (Thermo Fisher Scientific) per manufacturer’s recommendation. Media was changed 24 hours post transfection and 24 hours later, culture supernatants were collected spun down at 3000 rpm for 10 minutes, filtered and stored at -80°C. NIH 3T3 cells (0.2 × 10^6^ cells/well) were seeded in a 6-well plate and the next day were infected with luciferase reporter viruses normalized for equal amounts of p30 by western blots. Media was changed 24 hours post infection and luciferase levels were measured 48 hours post infection using Steady-Glo Luciferase Assay system (Promega) per the manufacturer’s recommendation and an automated plate reader, Biostack4 (Biotek) luminometer.

### Chloroquine and MG132 treatment

5 × 10^5^ 293T cells/well were seeded in a 6-well plate. Cells were co-transfected with 3 μg of an MLV infectious clone along with either 1 μg of E.V., mM1, or mM8. At 6 hours post transfection, fresh media ± 100 μM chloroquine (Sigma-Aldrich) or ± 16 μM MG132 (Sigma-Aldrich) was added. Cells were harvested and lysed in RIPA buffer 18 hours later followed by SDS-PAGE and western blot analysis.

### Statistical analysis

Statistical analyses were performed using GraphPad Prism software version 8.2. The statistical tests used to determine significance are described in the figure legends. A difference was considered to be significant for *P* values of <0.05.

## Supporting information

Suppemental Figure 3

## Acknowledgements

We thank Leonard “Pug” Evans, Susan Ross, Christine Kozak, Hans Orbea-Acha, Xinqi Liu, Benhur Lee, Amy Jacobs, Paul Bates and Ted Pierson for kindly providing us with reagents used for this project. We also thank Michael Battaglia and Sydney Herring for technical support with our cloning assays. The work was supported by National Institutes of Health grant R21-AI144147-01A1 and a Mathilde Krim Fellowship in Basic Biomedical Research Phase II, American Foundation for AIDS Research 109741-63-RKHF.

---

**Figure S1. Expression levels of mMARCH1, 2, 3, and 8 in the presence of IFN-β and MLV infection in EL4 and NIH3T3 cells.** Fold expression changes of mouse MARCH1, 2, 3 and 8 (mM1, 2, 3 and 8) relative to untreated cells and normalized to GAPDH in (A) EL4 mouse T cell line and (B) NIH3T3 mouse fibroblasts treated with 500 Units/mL of mouse IFN-β and harvested at 0, 4, 8, 16 and 24 hours post treatment. (C) Fold expression changes of mM1, 2, 3, and 8 relative to uninfected cells and normalized to GAPDH in (E) NIH3T3 cells at 4, 8, 16 and 24 hours post infection with MLV (5 MOI). (D) Purity of bone marrow derived macrophages (BMDMs) and (E) bone marrow derived dendritic cells (BMDCs) used in Fig 1 was confirmed by staining with either (D) the macrophage cell surface antibody CD11b-FITC or (E) the dendritic cell surface antibody CD11c-APC. Data shown represent averages of results from n=3 independent experiments. All results are presented as means ± standard error of the mean (SEM). Statistical analysis performed using unpaired t-test. ** = p ≤ 0.01, *** = p ≤ 0.001.

**Figure S2. Mouse MARCH1 and mouse MARCH8 localize in cellular membranes.** (A) Wild type (WT) mouse MARCH1 (mM1) and mM1 mutants (RING^mut^, ΔDIRT, TM1^mM3^, TM2^mM3^, TM1/2^mM3, 218^AAQL^221, 228^AARV^231^ and VQNC^mut^) localize in the plasma membrane of the transfected cells. 293T cells were transfected with mM1 constructs used in Fig 4 and 6. Cells were harvested and surface stained with anti-MARCH1 or rabbit IgG isotype control followed by anti-rabbit IgG (Alexa Fluor 647) and analyzed by flow cytometry. Histogram plots indicate percent of cells expressing mM1 on the cell membrane. (B) and (C) Wild type (WT) mouse MARCH8 (mM8) and mM8 mutants (RING^mut^, ΔDIRT, TM1^mM4^, TM2^mM4^, TM1/2^mM4, 218^AAQL^221, 228^AARV^231^ and VQNC^mut^) localize in the cellular membranes of the transfected cells. 293T cells were transfected with the various mM8 constructs used in Figure 4 and 6. Cells were harvested and integral membrane proteins were extracted using the MEM-PER plus membrane extraction kit (Thermo Scientific™) followed by western blots using anti-myc (mM8 detection) and anti-GAPDH (marker for purity of the membrane fractions). For empty vector (E.V) and mM8 WT transfection we provide the membrane fraction (M) and cellular fraction (C). Shown are the results of a single experiment (representative of two independent experiments).

**Figure S3. The VQNC motif does not affect mouse MARC1- and mouse MARCH8-mediated degradation of the retroviral envelope glycoprotein restriction.** 293T cells were co-transfected with an MLV infectious clone and either with mouse MARCH1 (mM1), mM8, mM1 with mutations in the VQNC domain (VQNC^mut^) or mM8 VQNC^mut^. Cells were harvested 48 hours post transfection and lysates were analyzed by immunoblotting using anti-MLV p30 (detects p65^Gag^), anti-MLV gp70, anti-MLV p15E, anti-myc (for detection of mM1 and 8) and anti-β-actin antibodies. Shown are the results of a single experiment (representative of three independent experiments).

**Figure S4. Mouse MARCH3 and mouse MARCH4 do not inhibit or physically interact with MLV p15E.** (A) Mouse MARCH4 does not target MLV envelope proteins for degradation. 293T cells were co-transfected with an MLV infectious clone and mouse MARCH4 (mM4). At 48 hours post transfection, cells and released virus in the culture media were harvested and the indicated proteins were analyzed by immunoblotting using anti-MLV p30 (detects p30 and p65^Gag^), anti-MLV gp70, anti-MLV p15E/p12E, anti-myc (for detection of mM4) and anti-β-actin antibodies. Shown are the results of a single experiment (representative of three independent experiments). (B) Mouse MARCH3 does not interact with MLV p15E. 293T cells were co-transfected with an MLV infectious clone and either (B) mM1 or mM3 and were harvested 48 hours post transfection. Cell lysates were immunoprecipitated with anti-myc (mM1 and mM3) followed by western blots probing with anti-myc (mM3) or anti-15E. Shown are the results of a single experiment (representative of three independent experiments).

**Figure S5. Viral envelope glycoproteins vary in their susceptibility to MARCH-mediated restriction.** (A) Lassa virus (LASV) mature envelope glycoprotein GP_2_, (B) Junín virus (JUNV) mature envelope glycoprotein GP_2_ (arrow), (C) Human Parainfluenza virus 1 (HPIV-1) hemagglutinin-neuraminidase (HN), (D) Measles virus (MV) hemagglutinin (H), (E) West Nile virus (WNV) envelope (E) and Crimean Congo hemorrhagic fever virus (CCHFV) mature envelope glycoprotein G_c_ protein levels in the presence of human MARCH1 (hM1), hM2, hM8 and empty vector (E.V.). 293T cells were co-transfected with either (A) LASV GPc, (B) JUNV GPc, (C) HPIV HN, (D) MV H, (E) WNV E or (F) CCHFV M segment along with either hM1, 2 8 or E.V. Cells were harvested 24 hours post transfection, lysed and analyzed by western blots using anti-V5 (for HPIV-1 HN and MV H detection), anti-FLAG (for LASV GP_2_ and JUNV GP_2_ detection), anti-Flavivirus E antibody (clone 4G2 for WNV E detection), anti-MARCH1, anti-MARCH2, anti-MARCH8 and anti-β-actin antibodies. Shown are the results of a single experiment (representative of three independent experiments).

## Notes

### Competing Interest Statement

The authors have declared no competing interest.

