## Supplementary figures and images for "Elucidating the antiviral mechanism of different MARCH factors"

### Suppemental Figure 3

#### Slide 1
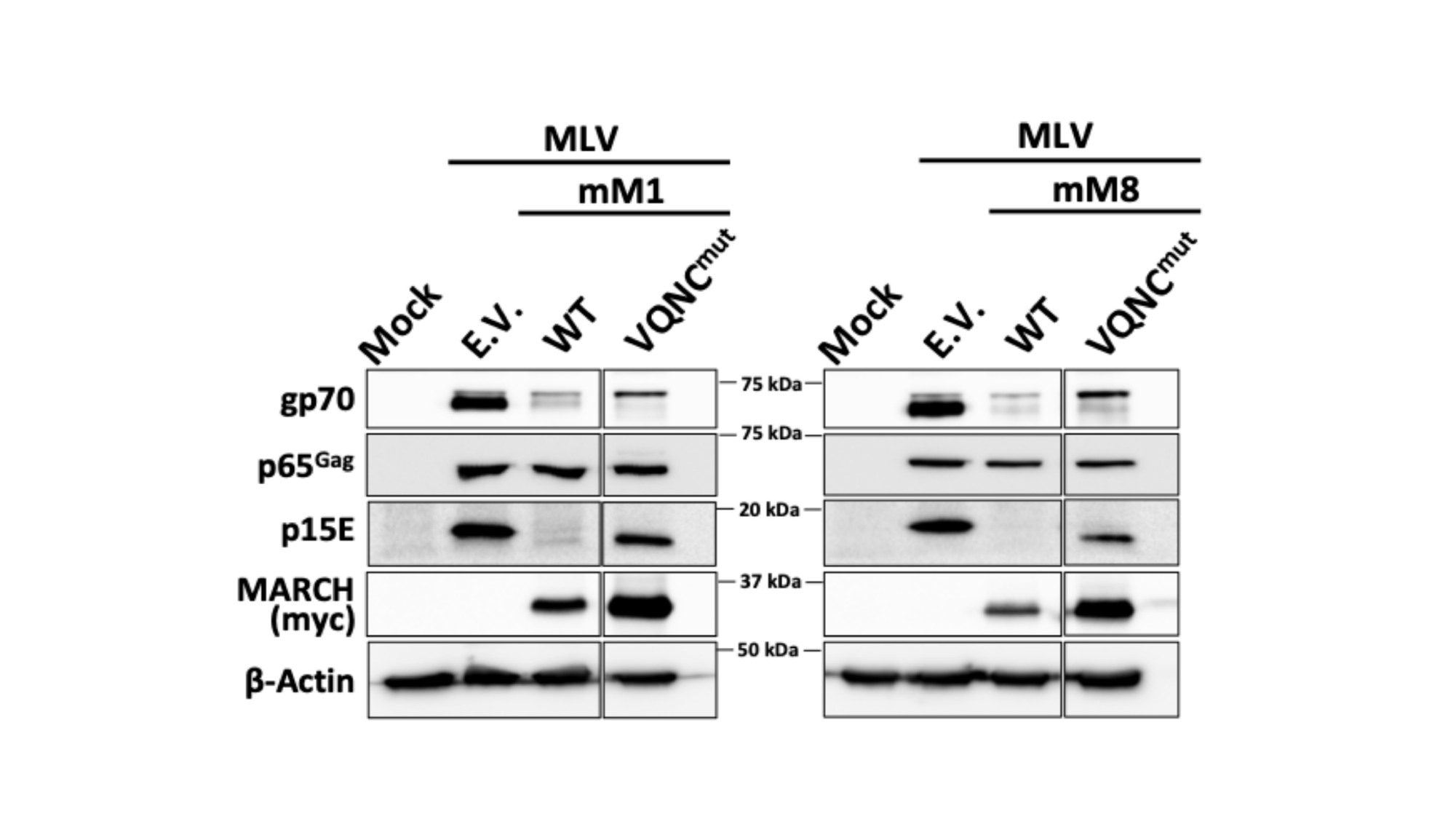

#

### Supplemental Figure 1

#### Slide 1
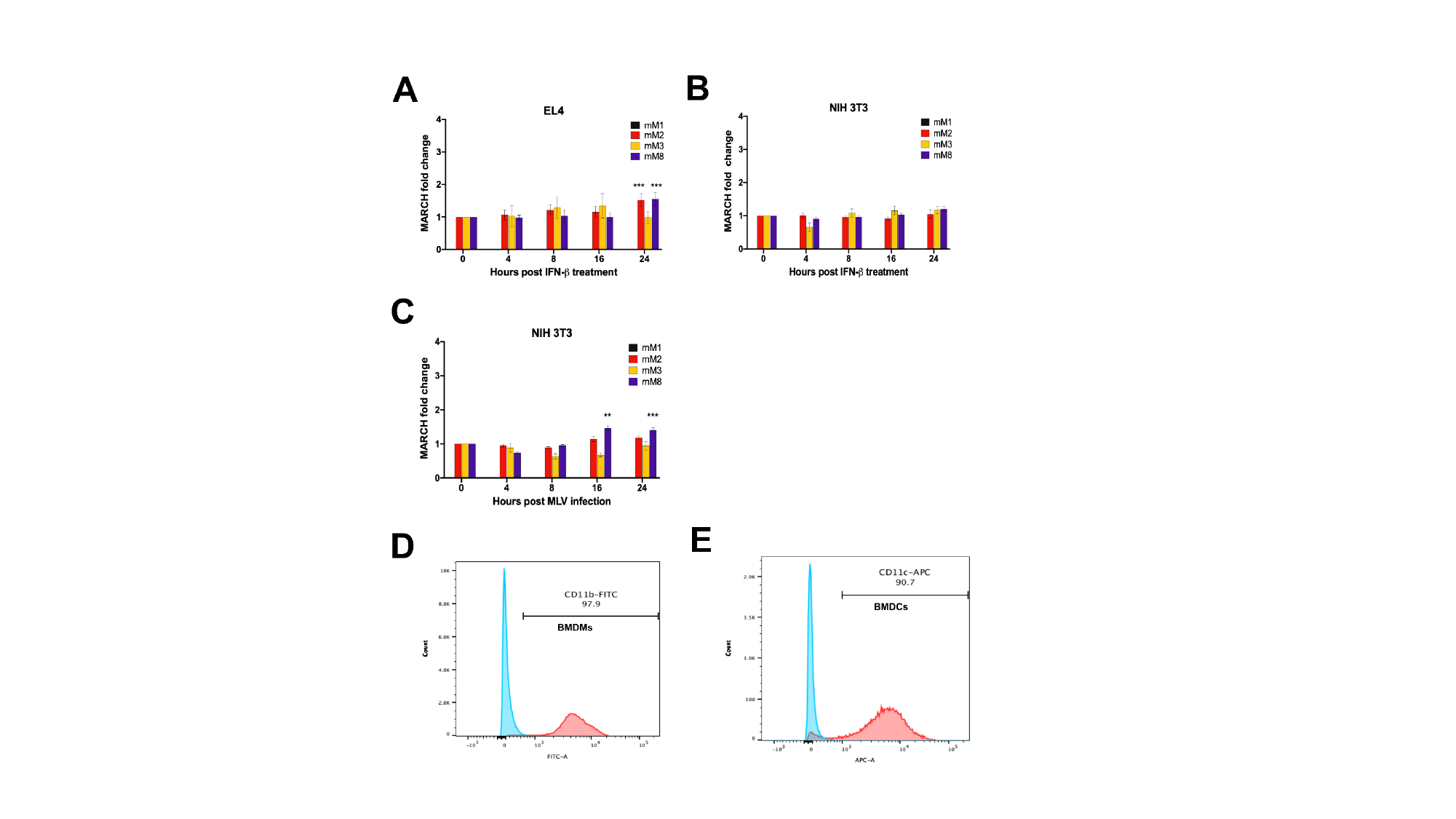

#

### Supplemental Figure 2

#### Slide 1
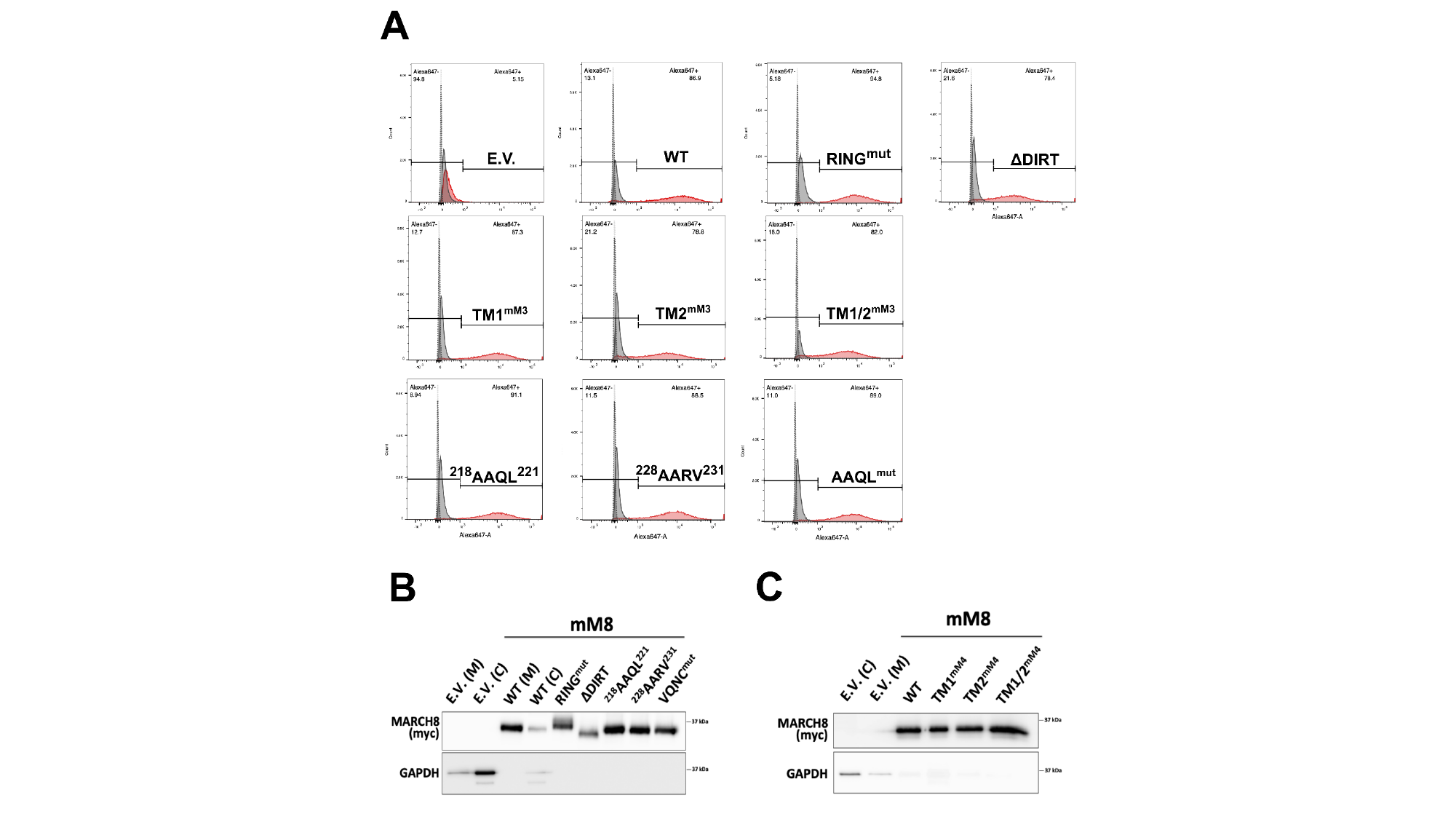

#

### Supplemental Figure 4

#### Slide 1
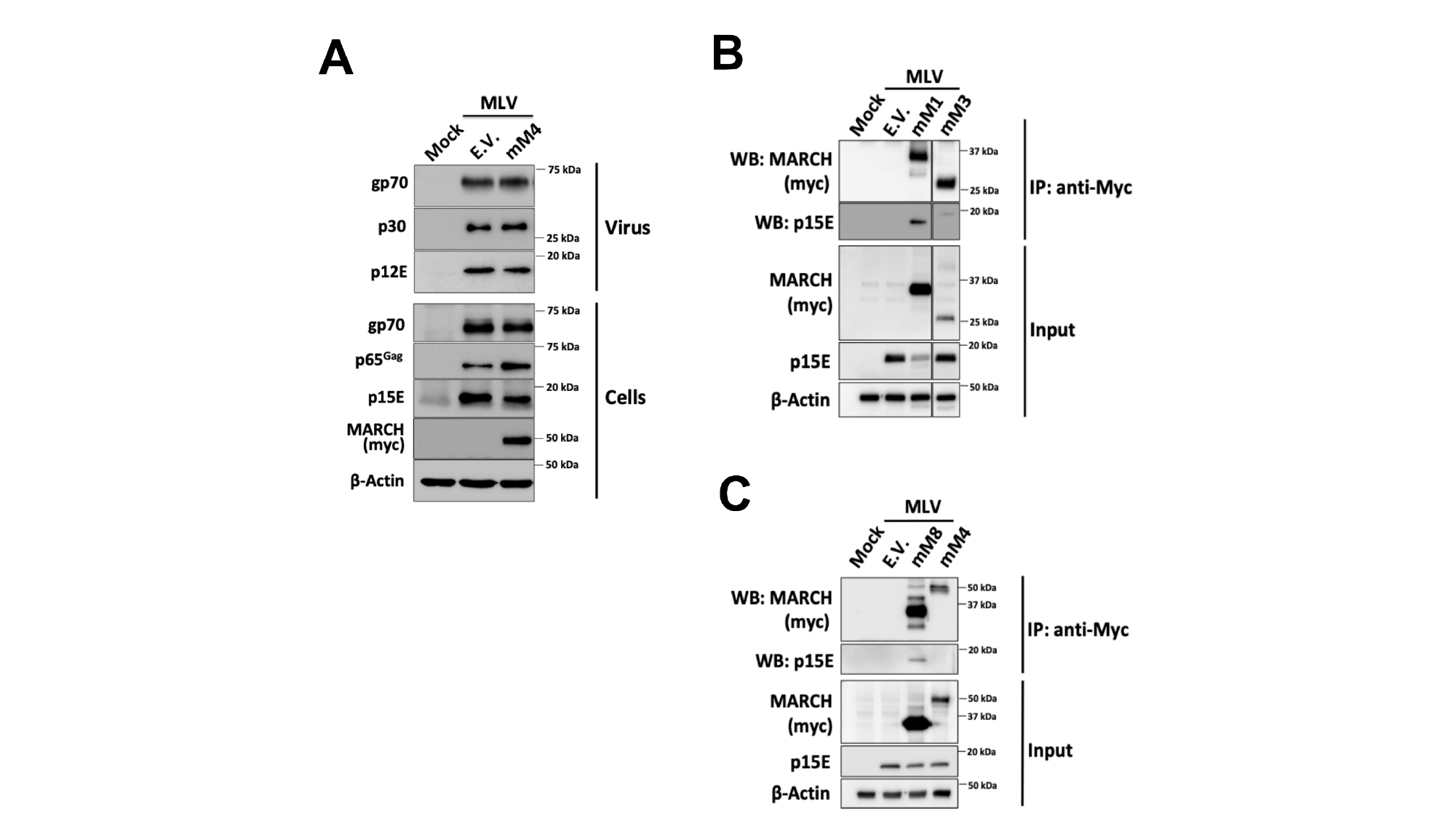

#

### Supplemental Figure 5

#### Slide 1
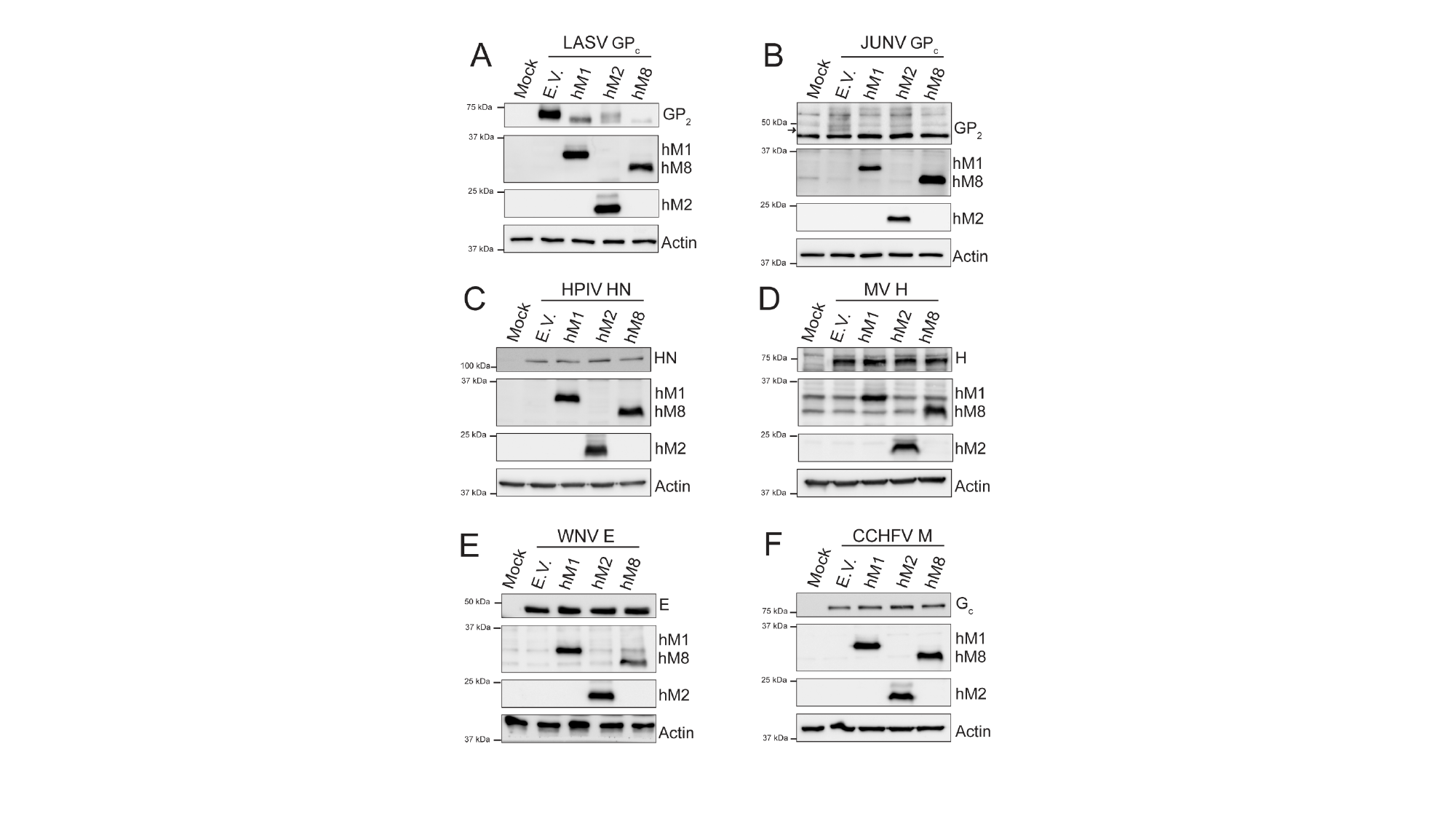

#
